# Fertilization offsets the maternal age effect on mouse oocytes provided the oviduct is not itself aged

**DOI:** 10.64898/2026.10.30.761756

**Authors:** Michele Boiani, Thomas Nolte, Julina Wiese, Georg Fuellen, Annalen Büchsenschütz, Saskia Winter, Hannes C.A. Drexler

**Affiliations:** Max Planck Institute for Molecular Biomedicine, Department of Cell & Tissue Dynamics, Röntgenstrasse 20, 48149 Münster, Germany; Rostock University Medical Center, Institute for Biostatistics and Informatics in Medicine and Aging Research (IBIMA), Ernst-Heydemann-Strasse 8, 18057 Rostock, Germany; Max Planck Institute for Molecular Biomedicine, Bioanalytical Mass Spectrometry Facility, Röntgenstrasse 20, 48149 Münster, Germany

**Keywords:** blastocyst, embryo, fertilization, maternal aging, oocyte, parthenogenesis, proteome, reproductive aging

## Abstract

Advanced maternal age compromises oocyte quality, and it is widely assumed that these deficits are transmitted to embryos. However, because most studies rely on *in vivo* conception, it has been difficult to separate the effects of oocyte aging from those of an aged fertilization environment. In aging mice, we observed that the fraction of ovulated oocytes forming implantation sites after *in vivo* conception fell ∼3.7-fold, raising the question of whether this was due to reduced fertilization, reduced embryo quality or reduced support from the aged reproductive tract. Therefore we fertilized oocytes of aged and young mice *in vitro* with a shared young-donor sperm preparation, holding the fertilization environment constant and removing paternal variability. Hallmarks of maternal age were largely offset following fertilization *in vitro*: aged oocytes produced blastocysts at rates comparable to young controls, with altered - but not deficient - cell-lineage composition and no detectable impairment in development to term (E18.5) after transfer to young oviducts. Proteomic analysis revealed that, unlike the transcriptomic signature, the oocyte’s proteomic age signature was not faithfully propagated to the blastocyst: for a subset of proteins, particularly those depleted in aged oocytes, the direction of the age effect reversed between oocyte and blastocyst. Comparison with parthenogenetically activated oocytes showed this divergence to be fertilization-dependent, implicating fertilization - rather than mere embryonic cleavage - as the driver of proteomic change. Stable-isotope labelling in live embryos showed that blastocysts from aged, fertilized oocytes both synthesized more new protein and retained less maternal protein than young counterparts; neither effect was detectable after parthenogenesis. Puromycin labelling, by contrast, showed a maternal-age-associated rise in bulk synthesis regardless of route. Together, these results indicate that the maternal-age effect on the oocyte proteome is labile rather than fixed and can be offset by fertilization itself, provided the maternal environment is not itself aged.

## Introduction

There is a notorious maternal age effect in mammalian reproduction, whereby the quantity and quality of oocytes decline as females get older. This effect is conserved across species, and has been modelled extensively in mice, with adverse age-associated outcomes broadly similar to those observed in humans (Flurkey et al., 2007; The Jackson Laboratories, 2017; Dalton-O’Reilly et al., 2023). It is commonly accepted that if maternally aged oocytes are of inferior cellular quality, then this state will propagate to embryos; with consequences that may extend beyond the embryo to the health of the resulting offspring. Propagation has been widely documented for chromosomal aneuploidies, which cannot easily be reversed once they have occurred, although the carrier cells can be segregated or eliminated (Bolton et al., 2016). However plausible, the propagation hypothesis relies on confirmatory narratives and does not take sufficient account of discordant observations, such as: 1) the reports in which embryos from aged mouse females developed to term without significant attrition when transplanted into younger recipients (Talbert and Krohn, 1966; Gosden, 1974; Suzuki et al., 1994; Woods et al., 2017; Zietek and Sampino, 2023; Gulzar et al., 2024; Cavanaugh et al., 2026); 2) the universally known fact that when aged females give birth, their offspring begin life biologically young, suggesting that somewhere during reproduction, aging was reversed (Chiavellini and Sebastiano, 2026).

Before we address the discordant observations, let us briefly recap the narrative of the maternal age effect on oocytes. Countless studies documented that the maternal age effect has three well-documented cytological-molecular hallmarks:

1. predisposition to aneuploidy due to reduced gene expression levels of chromosome cohesion factors, spindle assembly checkpoints, DNA damage response and subcortical maternal complex - SCMC – a cytoplasmic lattice required for the first embryonic cleavages (Hamatani et al., 2004; Steuerwald et al., 2007; Pan et al., 2008; Chiang et al., 2010; Grondahl et al., 2010; Titus et al., 2013; Lu et al., 2016; Barragan et al., 2017; Bebbere et al., 2020; Horta et al., 2020; Sharma et al., 2024);
2. impaired oxidative defense in conjunction with mitochondrial dysfunction (Kushnir et al., 2012; Zhang et al., 2014; Fragouli et al., 2015; Babayev et al., 2016; May-Panloup et al., 2019; Pasquariello et al., 2019; Morimoto et al., 2020; Hashimoto and Morimoto, 2022; Khan et al., 2023); and
3. compromised metabolism of transcriptional, post-transcriptional, translational and post-translational processes (Schwarzer et al., 2014; Duncan et al., 2017; Del Llano et al., 2020; Li et al., 2021; Huang et al., 2023; Galatidou et al., 2024; Dinh et al., 2025). However, these changes are not always observed in all studies. In the case of oocytes’ transcriptomes, for example, a meta-analysis of 12 studies found that independent corroboration of gene expression changes associated with age was actually limited to one or two studies (Zhang et al., 2025b).

Unlike oocytes, little is known about the maternal age effect on embryos. The reason is simple: because not all oocytes become fertilized and not all embryos progress to the next stage, the number of embryos available for study is more limited than the number of oocytes. The evidence supporting the propagation of defects is therefore fragmented into multiple studies sometimes small. The main burden transmitted to embryos is aneuploidy according to some investigators (Cozzolino et al., 2024), while other investigators contend that the decline in embryo quality occurs at various cellular levels, of which aneuploidy is just one among many (Reig et al., 2020). Indeed embryos of aged females in mice and humans have also fewer cells and more apoptosis at the blastocyst stage (Jurisicova et al., 1998; Velazquez et al., 2016), release inflammatory chemokines (Kawagoe et al., 2020), and express more of the ubiquitin-conjugating protein UBE2V1 (Li et al., 2025), which is suggestive of stress response tied to age-related proteostatic imbalance. “Omics” approaches allow to capture gene expression on a genome-wide scale rather than focusing on few hallmarks. Transcriptomic analyses revealed that maternal age is associated with large-scale transcript reductions at the blastocyst stage in humans (Kawai et al., 2018; McCallie et al., 2019) and also at pre-blastocyst stages in mice (Zhang et al., 2025c), as well as reduced protein machinery to perform RNA splicing at the 2-cell stage in mice (Li et al., 2021). It would be sensible to reach beyond the transcriptome, because the transcriptome is upstream in the gene expression cascade and its effects can therefore change on the way to protein. To date, proteome analysis has been conducted on human blastocoel fluid (Tedeschi et al., 2017), 2-cell stage mouse embryos (Li et al., 2021) and, more recently, 8-cell embryos obtained from clinically discarded tri-pronuclear zygotes of reproductively young and aged women (Zhang et al., 2025c). Thus, the proteomic dimension of the maternal age effect on embryos is under-studied.

We reasoned that the narrative of propagation of the maternal age effect typically rests on observations in which both the embryo and its environment were aged. Accordingly, we observed, in aging female mice, a 3.7-fold fall in the fraction of ovulated oocytes that implanted after *in vivo* conception. We therefore hypothesized that the maternal age effect might manifest differently if the age of the oocytes were decoupled from that of the mother’s body. To test this hypothesis, we subjected maternally young and aged mouse oocytes to parthenogenesis or *in vitro* fertilization (IVF) using a shared sperm preparation from young donor, thereby removing paternal variability and the aged reproductive tract. The embryos obtained in this way were compared in terms of developmental rates, as well as through analysis of the cellular and molecular composition of the blastocysts. In a nutshell, the defects of the aged oocytes were largely transmitted to parthenogenetic embryos while they were largely offset in the IVF embryos. Our results do not overturn the wider literature - nor could they, given that the proteomic analysis of blastocysts from maternally aged oocytes had not been carried out before. What these results do show is that, if the maternal environment is not itself aged, the impact of maternal age on embryos is less pronounced than previously thought, provided that the stimulus for development is the spermatozoon.

## Materials and methods

### Ethical permits for animal experiments

Mice were used for experiments according to two licenses issued by the Landesamt für Natur, Umwelt und Verbraucherschutz of the State of North Rhine-Westphalia, Germany, recently renamed as Landesamt für Verbraucherschutz und Ernährung NRW (LAVE) (license numbers LANUV AZ.81-02.04.2022.A376 “In-vitro und in-vivo-Markierung der Proteinsynthese in Mausembryonen”; LAVE AZ.2026-273 „Einfluss des maternalen Alters auf die Genexpression von Embryonen im Mausmodell“), in accordance with the guidelines laid down in the European Directive 2010/63/EU. We observed the ARRIVE guidelines to the extent applicable.

### Histological analysis of mouse ovaries

Ovaries were collected and immediately fixed in 4% paraformaldehyde (PFA) at 4 °C. Following fixation, tissues were dehydrated in a graded ethanol series (70%, 80%, 90%, 96%, and 100%), cleared in xylene, and infiltrated with molten paraffin using an automated tissue processor (Leica HistoCore, Leica Biosystems, Wetzlar, Germany). Processed tissues were embedded in paraffin blocks and sectioned at 8 µm thickness using a rotary microtome (HistoCore BIOCUT, Leica Biosystems, Wetzlar, Germany). Sections were mounted on glass slides and dried overnight at 37 °C. For histological staining and microscopy, sections were deparaffinized, rehydrated, and stained with hematoxylin and eosin according to standard histological protocols. Images were taken using a Nikon TE2000-U microscope fitted with a RisingCam 5109 camera.

### Crude chromosome counts

After removal of the zona pellucida with acidic Tyrode’s solution, oocytes were spread on slides according to the air-drying method (Tarkowski, 1966). Slides were stained with Hoechst 33342 (1 μg/mL) and scored on a Nikon TE2000 fluorescence microscope as described (Balbach et al., 2007).

### Metaphase II (MII) mouse oocytes collection

All mice used in this study were reared in-house at the MPI Münster. They were maintained in individually ventilated type 2 L cages (EHRET GmbH Life Science Solutions, 79111 Freiburg, Germany), with autoclaved Aspen wood as bedding material and a cardboard tube as enrichment, in groups of five females or individually as males. Access to water (acidified to pH 2.5) and food (Teklad 2020SX, ENVIGO RMS GmbH, Düsseldorf, Germany) was ad libitum. The animal room was maintained at a controlled temperature of 22 °C, a relative humidity of 55 %, and a 14/10 h light/dark photoperiod (light on at 6:00 a.m.). The hygiene status was monitored every 3 months, as recommended by Federation of European Laboratory Animal Science Associations (FELASA), and the sentinel mice were found free of pathogens that might have affected the results of this study. Ovulation was induced by i.p. injection of pregnant mare’s serum gonadotrophin (PMSG) (Pregmagon, IDT, 06861 Dessau-Roßlau, Germany) and hCG (Ovogest, MSD Tiergesundheit, Germany). B6C3F1 females aged 2 and 12-15 months were injected i.p. using a 27G needle, at 5 pm, with 10 I.U. PMSG and 10 I.U. hCG 72 h apart. The cumulus-oocyte complexes were recovered and dissociated in hyaluronidase to free the MII oocytes. Hyaluronidase (cat. no. 151271, ICN Biomedicals, USA; 50 I.U./mL in HCZB) was dissolved in Hepes-buffered Chatot, Ziomek and Bavister medium (HCZB) with bovine serum albumin (BSA; Probumin, Millipore, Kankakee, IL, USA) replaced by polyvinylpyrrolidone (PVP, 40 kDa; cat. no. 529504, Calbiochem, EMD Biosciences, La Jolla, CA, USA) 0.1% w/v.

### Proteome analysis of mouse oocytes in data-independent acquisition (DIA) mode

n=50 oocytes per sample were collected in triplicate for proteome analysis. The zonae were removed with acidic Tyrode solution. Oocyte samples (each in about 50µl LYSIS buffer) were prepared for MS analysis using the iST Kit (Preomics, Planegg/Martinsried, Germany) according to the manufactureŕs instructions. Nanoflow reversed-phase liquid chromatography was performed using a nanoElute 2 UHPLC system, online coupled to a timsTOF Pro2 mass spectrometer via a CaptiveSpray nano-electrospray source (all by Bruker Daltonics, Bremen, Germany). Peptides were separated on a 25 cm× 75 µm fused silica capillary column (ProtumLink, Jena, Germany), packed with 1.9 µm ReproSil-Pur 120 C18-AQ beads (Dr. Maisch, 72119 Ammerbuch-Entringen, Germany) using a 60 min linear gradient from 4 to 27% buffer B (99.9% acetonitrile, 0.1% formic acid; buffer A: 0.1% formic acid) at a flow rate of 300 nl/min. This was followed by a wash out phase for 7.5 min at 95% buffer B. Column temperature was maintained at 50°C. The mass spectrometer was operated in DIA-PASEF mode, using the standard DIA PASEF – long gradient MS-method. Signals covering a mass range from 100 to 1700 m/z and an ion mobility range from 1/K0 = 0.60 to 1.60 Vs/cm2 using a ramp time in the dual TIMS analyzer of 100 ms were recorded. The collision energy was lowered as a function of increasing ion mobility from 59 eV at 1/K= 1.60 Vs/cm2 to 20 eV at 1/K0 = 0.6 Vs/cm2. Accumulation time was set to 100 ms, source voltage to 1600V. Raw MS data were processed using DIA-NN (v. 2.3.2) in the library free search mode, using heuristic protein inference on the genes level. Neural networks were selected as machine learning-based search algorithm, matching between runs was enabled, and as quantification strategy QuantUMS was applied. For the murine database, entries were sourced from UniProt in January 2024. Carbamidomethylation on cysteine residues was set as fixed modification for the search, while oxidation at methionine and acetylation of protein N-termini were set as variable modifications. The maximum number of variable modifications was set to 1. Trypsin was defined as the digesting enzyme, allowing a maximum of 1 missed cleavages and a peptide length range of 7–30 amino acids. The maximum allowed mass deviation was 15 ppm for MS and 20 ppm for MS/MS scans. Charges above +5 were excluded. Protein groups were regarded as being unequivocally identified with a false discovery rate (FDR) of 1% for both the peptide and protein identifications. The oocyte proteomics data have been deposited in the ProteomeXchange Consortium via the PRIDE repository (Perez-Riverol et al., 2025) (http://proteomecentral.proteomexchange.org), with the dataset identifier PXD076504.

### Mouse embryo production *in vivo*

B6C3F1 females aged 2 or 12-15 months (young, aged) were hormonally stimulated as described, and mated to CD1 stud males aged 5-6 months. Seven days after successful mating, as indicated by the presence of a vaginal plug, the females were killed and implantation sites were scored.

### Mouse embryo production *in vitro*

Blastocysts were obtained from superovulated MII oocytes of B6C3F1 females aged 2 or 12-15 months (young, aged) either by *in vitro* fertilization (IVF) or parthenogenetic activation. Spermatozoa were isolated from the cauda epididymides of CD1 stud males aged 5-6 months. Epididymides were poked ≈20 times with 27G needles, placed on the bottom of a 2 mL plastic tube, and overlaid with 600 µL Whittingham medium containing 3% BSA. Swim-up was allowed to take place at 37°C under 6% CO_2_ for 30 minutes, after which the upper 100 µL layer were gently removed and placed as a drop in a 35 mm dish overlaid with preequilibrated mineral oil. Sperm concentration was adjusted to 2-2.5 million/mL and allowed to capacitate for 1 hour before adding the oocytes. Oocytes and spermatozoa were co-incubated at 37°C under 6% CO_2_ for 3-4 hours. Successful fertilization was indicated by the extrusion of a 2^nd^ polar body. Parthenogenetic activation was conducted in α-MEM medium lacking Calcium, containing 10 mM SrCl_2_ and 5 µM latrunculin B, for 6 hours. Successful activation was indicated by the formation of two pronuclei. After insemination or parthenogenetic activation, pronuclear oocytes were transferred to 500 µL of Potassium (K) simplex optimization medium (KSOM) in a four-well Nunc plate without oil overlay, at 37 °C under 6 % CO_2_ in air. KSOM was prepared in house as per original recipe and contained gentamicin (50 μg/mL), free amino-acids (aa) both essential and non-essential (hence called KSOM(aa)), 0.2% (w/v) BSA. For the labeling of embryonic protein synthesis using SILAC (Stable Isotope Labeling with/by Amino acids in cell Culture; Ong et al., 2002), KSOM(aa) medium was modified by replacing BSA with polyvinylpyrrolidone (40 kDa) and substituting the normal arginine and lysine with their isotopic variants Arg-10 (^13^C_6_H ^15^N_4_O_2_; cat. no. CNLM-539-H-PK, Cambridge Isotope Laboratories, Tewksbury, MA, USA) and Lys-8 (^13^C_6_H_14_^15^N_2_O_2_; cat. no. CNLM291-H-PK, Cambridge Isotope Laboratories), as described (Israel et al., 2023). Zygotes were cultured in KSOM(aa) at 37°C under 6% CO_2_. At 96 hours post-IVF the blastocysts were collected and processed further.

### Analysis of cell lineage allocation at the blastocyst stage

Blastocysts were fixed with 3.7 % formaldehyde in 1x PBS for 20 min at 4 °C, permeabilized with 0.1% Triton X-100 in 1x PBS for 20 min at 4 °C, and then blocked with 2 % BSA, 2 % glycine, 5 % donkey serum, 0.1 % Tween20 in 1x PBS overnight at 4 °C. Fixed blastocysts were subjected to triple immunostaining followed by confocal microscopy imaging to identify and map the different cell lineages, as described previously (Schwarzer et al., 2012). The following primary antibodies were applied simultaneously to the specimens overnight at 4 °C for analysis of cell lineage allocation: anti-Cdx2 mouse IgG1 (Emergo Europe, The Hague, Netherlands, cat. no. CDX2-88), anti-Nanog rabbit IgG (Cosmo Bio, Tokyo, Japan, cat. no. REC-RCAB0002P-F) and anti-Sox17 goat IgG (R&D Systems, cat. no. AF1924) in dilutions of 1:200, 1:2,000 and 1:100, respectively. Appropriate Alexa Fluor-tagged secondary antibodies (Invitrogen) were matched to the primaries and incubated for 1–2 h at room temperature. For imaging, embryos were placed in 5 µl drops of PBS on a 50-mm thin-bottom plastic dish (Lumox hydrophilic dish, Greiner Bio-One, Frickenhausen, Germany) and overlaid with mineral oil (cat. no. M8410, Sigma-Aldrich Chemie GmbH, 82024 Taufkirchen, Germany). Images were captured using a 20x CFI Plan Apochromat VC objective on an inverted motorized Nikon TiE2000 microscope fitted with an Andor Dragonfly 502 spinning disc confocal unit Scanning System. Optical sections per embryo were captured using a high resolution (2048 x 2048 pixel) sCMOS camera. Maximum projections were analyzed with Fiji (Schindelin et al., 2012).

### Transfer of mouse IVF embryos to pseudo-pregnant females

Embryos were surgically transferred in groups of ten to the genital tract of pseudo-pregnant ≈9 week-old CD1 females weighing ≈30 grams. We transferred 4-cell embryos into one oviduct of females that had received the copulation plug from vasectomized CD1 males on the same day as the embryo transfer after three days of caging them together. Prior to surgery, pseudo-pregnant females were anesthetized with Ketamine (80 mg/kg body weight)/Xylazin (16 mg/kg)/Tramadol (15 mg/kg) in PBS, delivered i.p. The anesthetized mouse was placed on a heated pad set to 37 °C. Access to the oviduct was achieved via a dorsal laparotomy, involving a midline incision of the skin and a lateral incision of the peritoneum. The ovary and the oviduct were gently exposed, the ovarian bursa was torn, and the embryos were deposited into the infundibulum using a glass pipette with flame-polished tip. After that the ovary and the oviduct were returned to the original location, and the skin wound was closed with Michel Suture Clips (7.5 x 1.75 mm; Fine Science Tools, Germany). The mouse was placed in a cage heated to 30 °C under an infrared lamp. Post-surgical pain was alleviated by providing the animals with Tramadol in drinking water (0.5 mg/mL). Pregnancies were evaluated by Caesarean section just prior to term (embryonic day 18.5).

### Transcriptome analysis of mouse blastocysts

Total RNA was extracted and purified using Quick-RNA MicroPrep (cat. no. R1051, Zymo Research Europe GmbH). The library preparation of the total RNA was performed with the Watchmaker mRNA Library Prep Kit (7K0078-024), which does not use poly(A) tail selection, according to the manufacturer’s instructions. Paired end sequencing with a read length of 111bp was performed on NovaSeq® X Plus System using the corresponding NovaSeq® X Series 1.5B Reagent Kit (200 Cyc). Total RNA integrity and quality of the library were assessed using a TapeStation4200 (Agilent, Santa Clara, CA, USA). On average, the libraries contained 13.6±2.1 million 111-base reads. Using a molecular barcode, the samples were demultiplexed (DRAGEN BCL Convert 4.3.13) to fastq data and quality controlled (FastQC version 0.12.1; https://www.bioinformatics.babraham.ac.uk/projects/fastqc/). Trimmomatic (Bolger et al., 2014, version 0.39) was used for adapter trimming and read filtering. Reads were aligned to the reference genome (Mus musculus Ensembl GRCm39) using Hisat2 (Kim et al., 2019, version: 2.2.1). The aligned reads were sorted using samtools (Li et al., 2009). Sorted and aligned reads were counted into genes using HTSeq framework (Anders et al., 2015, version 2.0.3), then these counts were used to calculate the FPKM to normalize for sequencing depth and gene length. StringTie version 3.0.0 was used to calculate the potential transcripts. FPKM values for Ensembl gene identifiers corresponding to the same gene symbol were averaged. The blastocyst transcriptomics data have been deposited in the ArrayExpress collection in BioStudies under accession numbers E-MTAB-16116.

### Low-input proteome analysis of metabolically labeled blastocysts in data-independent acquisition (DIA) mode

The blastocysts were processed for proteomics largely following the method of Zhang and colleagues (Zhang et al., 2025a) with some modifications. Briefly, blastocysts were deprived of the zona pellucida (acidic Tyrode solution) and collected in pools of 3 blastocysts per sample in 8 μL lysis and digestion buffer deposited at the bottom of conical 250-µL autosampler glass inserts (Macherey Nagel, cat. no. 702716). The composition of the buffer was: 0.2 % n-Dodecyl-β-D-Maltoside, 100 mM HEPES, and 20 ng/μL sequencing grade modified Trypsin (Promega). Inserts were placed in autosampler vials, sealed, and incubated for 2 h at 37°C in a water bath to complete tryptic digestion. Samples were then stored at −20°C without further cleanup or desalting until LC–MS analysis. Nano liquid chromatography-tandem mass spectrometry (nanoLC-MS/MS) analysis was performed on a timsTOF pro mass spectrometer (Bruker Daltonics, Bremen, Germany) coupled to a nanoElute 2 ultra-high-performance liquid chromatography (UHPLC) system via a CaptiveSpray nano-electrospray ion source. Peptide samples were separated using a fused silica capillary column with integrated emitter tip packed with 1.9 μm / 120 Å C18 resin (25 cm L x 360 µm OD x 75 µm ID; ProtumLink core, Jena, Germany). Chromatography was accomplished using a multi-linear 60 min gradient from 4 % - 20 % buffer B (0.1 % formic acid in acetonitrile) over 42 min, from 20 % – 37 % buffer B over 6 min and from 37 % - 85 % buffer B in 4 min, followed by a 8 min wash-out at 85 % buffer B (buffer A: 0.1 % formic acid in water; flow rate: 300nl/min; column temperature: 50 °C). The mass spectrometer was operated in DIA-PASEF mode using the vendor “DIA-PASEF long gradient” method. MS signals were acquired over m/z 100–1700 and 1/K0 0.60–1.60 V·s/cm² with a dual TIMS ramp time of 100 ms. Collision energy was stepped as a function of ion mobility from 59 eV at 1/K0 = 1.60 V·s/cm² to 20 eV at 1/K0 = 0.60 V·s/cm². The accumulation time was 100 ms, and the source voltage was set to 1600 V. Raw data were processed with DIA-NN v2.3.2 in library-free mode with heuristic protein inference at the gene level. Neural networks were used as the search engine, match-between-runs was enabled, and QuantUMS quantification was applied. The murine protein database was derived from UniProt (April 2024). Heavy and light amino acids K and R were specified as fixed modification using the additional option commands --fixed-mod SILAC,0.0,KR,label, --lib-fixed-mod SILAC, --channels SILAC,L,KR,0:0; SILAC,H,KR,8.014199:10.008269, --peak translation, --original-mods and –channel-run-norm. Methionine oxidation and protein N-terminal acetylation were set as variable modifications (maximum three variable modifications per peptide). Trypsin was specified as the cleaving enzyme allowing one missed cleavage and a peptide length range of 7–35 amino acids. The maximum mass deviation was 15 ppm for precursors and 20 ppm for fragments; precursor charges >+5 were excluded. Protein groups were reported at 1% FDR at both peptide and protein levels. The blastocyst proteomics data have been deposited in the ProteomeXchange Consortium via the PRIDE repository (Perez-Riverol et al., 2025) (http://proteomecentral.proteomexchange.org), with the dataset identifier PXD076689 (fertilized blastocysts) and PXD082343 (parthenogenetic blastocysts).

### Cluster analysis and over-representation analysis of differentially expressed proteins

Proteome data analysis was performed in-house using the output of the DIA-NN program, exported to Microsoft Excel (Microsoft Deutschland GmbH, 80807 München, Germany) format and imported in JMP Pro v. 18 (JMP Austria, Germany & Switzerland, 69188 Heidelberg, Germany). We considered the proteins that were present in all samples of all replicates, and their values were Z-normalized for downstream cluster analysis. Over-representation analysis was performed with the WEB-based GEne SeT AnaLysis Toolkit (WebGestalt, version 2024; Elizarraras et al., 2024) against the GO ‘cellular component’ ontology, using the protein-coding genome as reference set. Because a detected proteome is not a random draw from the genome, we repeated every analysis using as reference the set of proteins detected in the corresponding experiment (3611 in oocytes, 3793 in IVF blastocysts, 2757 in parthenogenetic blastocysts).

### Cross-stage redistribution of age-affected proteins

For each protein quantified at both ages, the effect of maternal age was determined separately in the oocyte and in the blastocyst, and each protein was assigned to one of three categories per age - “DOWN”, “STABLE” or “UP” in reproductively aged (12-15 months, aged, for short) relative to young based on the z-scored abundances, with clusters labelled by their mean aged-young difference (k-means clustering, k = 3). Combining the oocyte and blastocyst assignments partitioned the co-detected proteins into 3 × 3 = 9 mutually exclusive flux categories (DOWN→DOWN, DOWN→STABLE, DOWN→UP, STABLE→DOWN, STABLE→STABLE, STABLE→UP, UP→DOWN, UP→STABLE, UP→UP), where the first term denotes the oocyte category and the second the blastocyst category. The fluxes were rendered graphically using Sankey diagrams. The number of proteins in each category of flux was tabulated in a 3 × 3 contingency table. We first tested this table for departure from independence using χ² test, which compares the observed category counts with those expected if the events were independent. Because the χ² independence null assumes that each protein measurement is an independent observation (an assumption violated by the correlated noise structure of proteomic data, and one that can be further inflated when proteins selected at the extreme of one dataset regress toward the mean on independent re-measurement in the other), we validated the result against an empirical null obtained by label permutation. For each of 1000 permutations, the young/aged sample labels were randomly reassigned within each stage independently (preserving group sizes: 3 versus 3 oocyte and 5 versus 5 blastocyst replicates), the entire pipeline was re-run on the permuted data (per-protein age effects, three-way classification, and 3 × 3 cross-tabulation), and the χ² statistic, the count in each category of flux, and the correlation between the oocyte and blastocyst age effects across co-detected proteins were recorded. Empirical 2-sided p-values were computed as the proportion of permutations yielding a value at least as extreme as the observed one, with the true labeling excluded from the null. Analyses were performed in Python (v3) using NumPy, pandas and SciPy. To test directly whether the oocyte–blastocyst relationship differed between fertilization and parthenogenesis, the two Spearman correlations were compared with Fisher’s z transformation, applying the variance inflation factor of 1.06 appropriate to ρ.

### O-propargyl-puromycin (OPP) assay for imaging of *de novo* protein synthesis

Using the original method with modifications (Liu et al., 2012), embryos were incubated with 20 µM OPP dissolved in KSOM(aa) medium for 3 h. In an initial control experiment, embryos were precultured in 50 µg/ml cycloheximide for 2 h before applying OPP, to confirm that the signal produced by OPP incorporation was genuine. Treated and control embryos were fixed with 3.7 % formaldehyde for 30 min and permeabilized with 0.5 % Triton X-100 for 20 min. For detection of the OPP, embryos were incubated with the CLICK reaction cocktail containing the red-fluorescent AZDye555-Azide (prepared according to the protocol from Jena Bioscience) for 60 min at RT, protected from light. Control embryos were subjected to the detection reaction without prior OPP incubation. Cell nuclei were stained with Hoechst 33342. All reagents were then washed off in a large excess volume of PBS. For imaging, embryos were pipetted into a 3 µl drop of PBS overlaid with oil on a glass-bottomed petri dish. Images were taken on a Nikon TE2000-U microscope fitted with a sCMOS camera. The brightfield microscopy and DNA staining images were captured with an exposure time of 10 ms. The fluorescence signals of the immunolabeled samples were detected with an exposure time of 50 ms. Fluorescence intensities were measured using Image-J, and the intensity of the detection control was subtracted from those of the OPP.

### Statistical analysis of developmental rates and gene expression levels

The two-sided Wilcoxon rank-sum (Mann–Whitney) test was used to calculate p values unless otherwise stated. With n = 3 per group the smallest attainable two-sided rank-sum p is 0.10, so parthenogenetic comparisons at n = 3 (SILAC/DIA channels) can exclude a large effect but not any effect; p = 1.00 in those panels indicates no rank separation, not equivalence. Exceptions were the refinement of the directional clustering analysis (t-test) and the refinement of cross-stage redistribution of age-affected proteins (χ² test with a permutation null). Calculations were performed using the statistical program JMP v.18 (JMP Austria, Germany & Switzerland, 69188 Heidelberg, Germany). No multiplicity correction was applied unless otherwise stated.

## Results

### Maternally aged mouse oocytes are fewer and larger, and their proteome is directionally redistributed

We ensured, before embarking in the analysis of embryos, that features expected of the maternal age effect were present in the starting biologic material, as modeled in the B6C3F1 mouse strain. This strain was chosen for the uniformity of the oocytes (B6C3F1 being an F1 hybrid of inbred strains) and for its robust developmental competence. We surveyed ovulated oocyte yield across ages from 2 to 29 months. We focused the young and aged arms of the study on 2 and 12–15 months, roughly equivalent to women in their 20s and 40s, respectively. The aged ovaries were manifestly depleted of follicles (Figure 1A) and the number of oocytes ovulated in stimulated ovarian cycles had already fallen by ≈70% compared to young age (median 32 vs 9; Figure 1B; Supplementary Table S1), while nearly all the aged females were still alive at 12–15 months (Yamamoto et al., 1993), thereby ruling out a survivor bias in this study. Furthermore, and as expected, crude aneuploidies had increased at 12–15 compared to 2 months (Figure 1C). Although the clutches of ovulated oocytes were smaller at 12–15 months, the oocytes themselves were slightly larger than the young counterparts, as inferred from the vitelline diameter (median 77.7 vs 75.3 µm; p < 0.0001, Wilcoxon; Figure 1D,D’; Supplementary Table S2).

**Figure 1.**
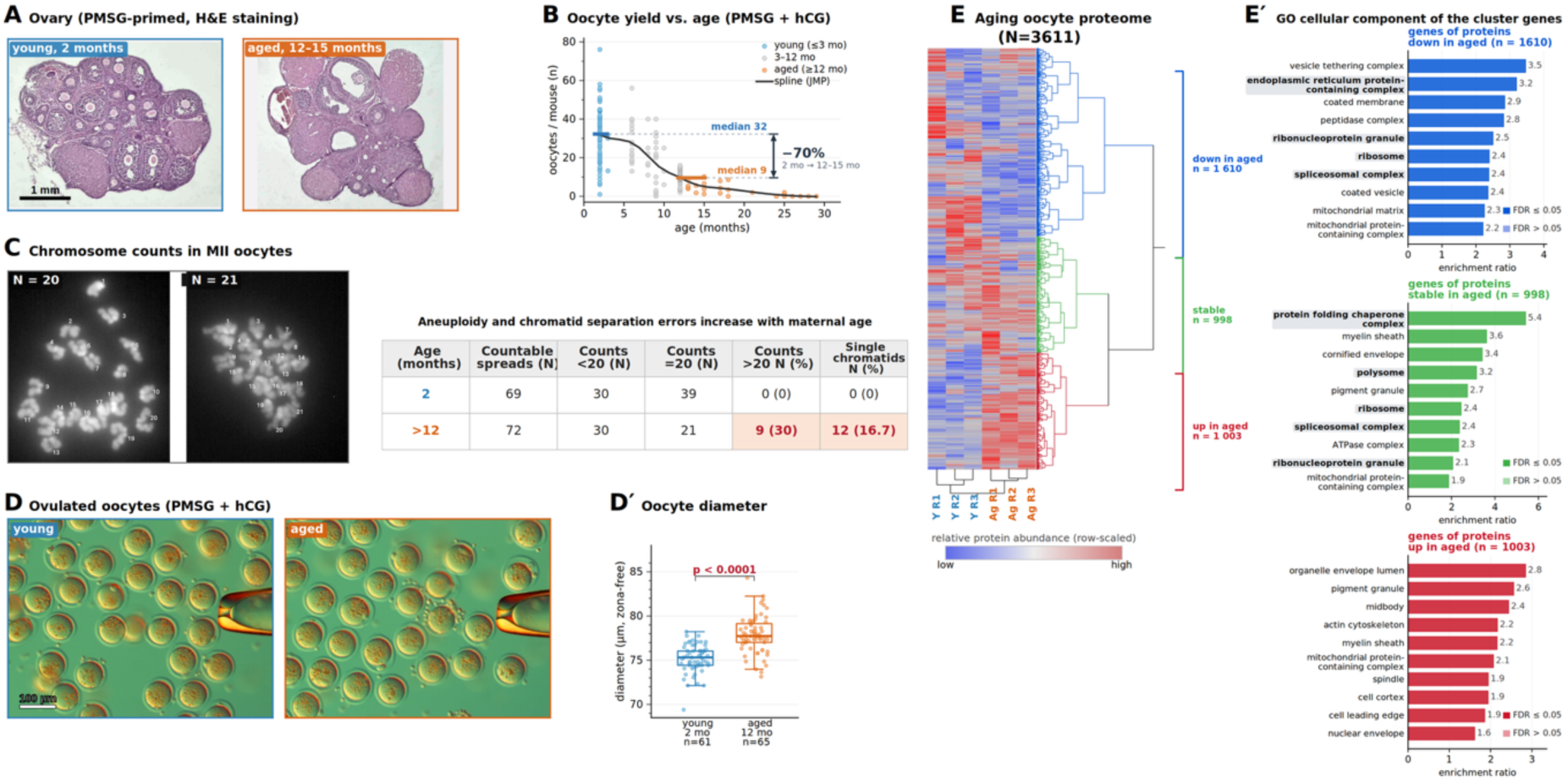
The starting biologic material shows the features expected of the maternal age effect. Young = 2 months, aged = 12–15 months (B6C3F1) throughout. (A) Ovarian sections (H&E, 8 µm) showing follicle depletion with age; images cropped and white-balanced separately, not at a common scale. Scale bar = 1 mm. (B) Oocytes recovered after superovulation vs maternal age (dot = one female; curve = spline); yield declines with age. (C) Chromosome counts in MII chromosome preparation. Spreads with <20 chromosomes (30/69 young, 30/72 aged) reflect loss during spreading and were equally frequent, so the comparison rests on hyperploidy and single chromatids: among spreads with ≥20 chromosomes, both were confined to the aged group (hyperploidy 9/30 vs 0/39; single chromatids 12 vs 0; Fisher’s exact p = 3 × 10⁻⁴ each). (D) Brightfield oocytes from a young and an aged female superovulated the same day (isolated 14 h post-hCG, hyaluronidase-treated). Scale bar = 100 µm. (Dʹ) Equatorial vitellus diameter (Fiji): aged oocytes are larger (median 77.7 vs 75.3 µm, ≈10% volume; n = 61 young, 65 aged; Wilcoxon). Box plots overlaid with individual data points show the interquartile range, the central line the median and the whiskers the data range excluding outliers. (E) Two-way clustering of the oocyte proteome (3 young, 3 aged; 50 MII oocytes each): proteins quantified in all six samples were z-normalized and grouped by k-means (k = 3, rows); samples (columns) clustered hierarchically and assigned correctly by age. Color = row-scaled abundance (blue = low, red = high). (Eʹ) Over-representation analysis per cluster (WebGestalt, cellular component; reference = protein-coding genome), as enrichment ratio; all displayed terms pass FDR ≤ 0.05, so the open/shaded-bar convention for FDR > 0.05 shown in the key does not occur in this plot. GO CC terms related to proteostasis and mRNA splicing are written bolded on gray background. Abbreviations: PMSG, pregnant mare’s serum gonadotropin; hCG, human chorionic gonadotropin; H&E, Hematoxylin-Eosin; MII, metaphase II; Y, young; Ag, aged. **Alt text.** Characterization of ovaries and oocytes from young (2-month) and aged (12–15-month) B6C3F1 mice. Panel A, stained ovarian sections showing fewer follicles with age; B, plot of oocyte number falling with maternal age; C, representative images and a table of chromosome counts in MII chromosome preparations; D, brightfield images of young and aged oocytes with Dʹ box plots showing aged oocytes are slightly larger; E, a two-way clustered heat map of the oocyte proteome, showing the three directional changes that occur with maternal aging; E‘, top-10 enriched GO CC of the proteins that increase, keep stable or decrease with maternal aging.

Ovulated oocytes were collected in triplicate from young and aged females in parallel, after ovarian stimulation, and subjected to proteome analysis by LC-MS/MS. Of 4889 proteins detected overall, 3611 were present in all 6 samples (range 4042-4256 per sample; Supplementary Table S3). Since there are only three ways in which maternal age can influence protein abundance (DOWN, STABLE and UP with aging), we applied k-means (k = 3) clustering on the Z-normalized values of the 3611 proteins (rows) and hierarchical clustering on the 6 samples (columns). Sample clustering correctly assigned the samples to the two age groups (young, aged) (Figure 1E). Protein clustering produced three clusters of different sizes, that is, DOWN in aged, n=1610; STABLE, n=998; UP in aged, n=1003 (Figure 1E). It should be noted that this clustering is directionally sound but statistically permissive: cluster memberships are directional assignments, and only a subset of the proteins in each cluster attained statistical significance (DOWN in aged, n=282 of 1610; STABLE, n=73 of 998; UP in aged, n=608 of 1003; t-test, uncorrected p < 0.05). Using the protein-coding genome as a reference, we performed over-representation analysis (Webgestalt) of the directional clusters in the gene ontology ‘cellular component’ (GO CC), chosen owing to its direct link to observable, physical structures of the cell. GO CC analysis returned terms related to ribosomes and mRNA splicing among the proteins DOWN with maternal age (FDR ≤ 0.05), consistent with the literature on the maternal age effect on oocytes (e.g. Galatidou et al., 2024), while terms related to mitochondria were not cluster-specific, appearing among the top-10 terms of all three directional clusters (Figure 1E’). Because the choice of reference set influences over-representation analysis (Wijesooriya et al., 2022), we repeated the GO CC analyses using the detected proteome as reference (Supplementary Figure S1). Fewer terms passed FDR ≤ 0.05, as expected given the smaller reference set. Term identity was broadly preserved, with three to five of the ten top terms shared per cluster. In particular, and in line with the oocyte literature (Introduction), protein synthesis continued to be downregulated with maternal aging, as inferred from the GO CC term “endoplasmic reticulum protein-containing complex” (which is another name for rough endoplasmic reticulum - the site of protein synthesis), whilst actin continued to be upregulated in oocytes as maternal age increased. Unexpectedly, among the maternal-effect factors that build the SCMC (Li et al., 2008; Bebbere et al., 2021), none was depleted with age: the core components (NLRP5, OOEP, TLE6, KHDC3, PADI6) partitioned into the STABLE cluster (Figure 1E), and several SCMC and SCMC-associated proteins were more abundant in aged oocytes (e.g. PADI6 +22%, p = 0.10; NLRP2 +21%, p = 0.10; ZBED3, p = 0.10; Wilcoxon; Supplementary Table S3). Maternally aged oocytes therefore enter embryogenesis with their SCMC almost intact - a starting point we will revisit when examining the fate of these proteins in embryos.

Collectively, these data document that key features expected of the maternal age effect (Introduction) were present in the biological starting material of this study (e.g. ovaries were depleted of follicles, ovulated clutch size was reduced by 70%, aneuploidy was increased, ribosome/splicing terms in GO CC analysis were affected in aging). The effect of maternal age is not uniform; that is, not all proteins are affected equally, as we had proposed in our previous study (Schwarzer et al., 2014). Therefore, we used this confirmed material to test whether the hallmarks of maternally aged oocytes are passed on to the embryos.

### Embryos of maternally aged mouse oocytes vanish after fertilization *in vivo*

Having established that aged B6C3F1 females ovulate fewer oocytes and that these exhibit proteome alterations, we asked what becomes of those oocytes once they are fertilized in their own natural environment. In two independent cohorts, young and aged females were superovulated as described, followed by the counting of oocytes in the oviductal ampulla or by mating. Implantation sites were scored in the uterus on day 7 after mating (Figure 2A), before any overcrowding and animal welfare issues could arise. Compared to the reduced number of oocytes superovulated in aged compared to young females (39 → 8, -79%; Figure 2B), a substantially smaller proportion of these oocytes formed implantation sites in aged females (18 → 1, -94%; Figure 2C) (Supplementary Table S4). In other words, there was a 3.7-fold fall in the fraction of ovulated oocytes that implanted, in aged females compared to young females. This is the point at which other investigators would probably be satisfied, arguing that the figures add up: there are fewer implantations than expected, because the maternally aged oocytes were of lower quality. To us, things seem less straightforward.

**Figure 2.**
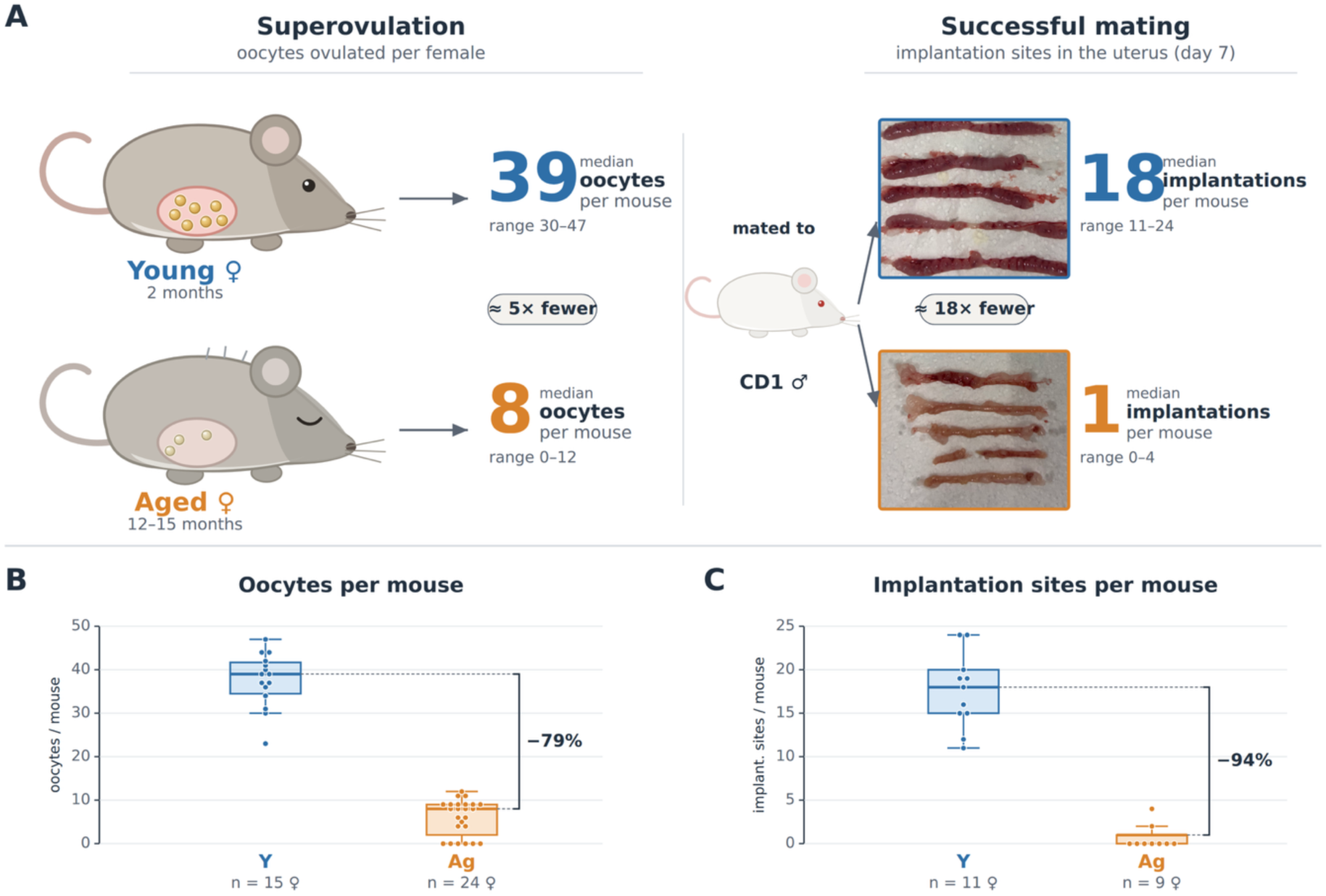
More pronounced loss of embryos in aged females than in younger females. **(A)** Schematic of the two *in vivo* readouts. Oocytes superovulated per female after hormonal stimulation, in young and aged females (♀). These were mated to CD1 males (♂), and following successful mating, implantation sites were counted in the uterus at day 7 of pregnancy; the photographs show dissected day-7 uteri (five specimens per cohort). Large numerals give the median per female (oocytes, 39 vs 8, -79%; implantation sites, 18 vs 1, -94%; young vs aged respectively) with the range beneath; rounded labels indicate the approximate fold reduction with maternal age (≈5× fewer oocytes, ≈18× fewer implantation sites). (B, C) Oocytes per female after superovulation (B) and implantation sites per female after successful mating (C) in young (Y) and aged (Ag) mice. Boxes show the interquartile range, the central line the median and the whiskers the data range excluding outliers; each dot is one female. n, number of females, is given beneath each group. Young = 2 months, aged = 12–15 months (B6C3F1) throughout; ♀, female; ♂, male. **Alt text.** Reproductive output of young (2-month) versus aged (12–15-month) female mice (♀). Panel A, schematic: aged females ovulate fewer oocytes (median 8 vs 39) and, after successful mating to CD1 males, have far fewer implantation sites as scored on day-7 (median 1 vs 18); the photographs show uterine horns rich in decidua in young and nearly empty in aged females. In panels B and C, box-and-whisker plots overlaid with individual data points show the age-related fall in oocytes (B) and implantation sites (C); n is the number of females.

Three explanations can account for the gap between oocyte numbers (Figure 2B) and implantations numbers (Figure 2C): the aged oocytes may fail to be fertilized, or they may be fertilized but experience attrition during preimplantation due to the lower oocyte quality, or they may not be adequately supported by the aged reproductive tract; a combination of these factors is also possible. The *in vivo* setting cannot distinguish between these possibilities, because fertilization and early development take place hidden within the oviduct and uterus - compartments that are themselves subject to maternal aging. To separate a failure of fertilization from a loss after fertilization, and to isolate the oocyte from its aged environment, it is necessary to abandon the *in vivo* setting and reconstruct fertilization *in vitro*, where each step can be scored directly. We therefore turned to *in vitro* fertilization (IVF).

### Maternally aged mouse oocytes fertilized *in vitro* yield blastocysts without the expected functional deficits

Embryos were produced by IVF of B6C3F1 oocytes, using spermatozoa of the same CD1 male for the oocytes of both the aged and young females, thereby equalizing the paternal effect and the fertilization environment (Figure 3A). Oocytes that extruded a 2^nd^ polar body were considered fertilized, and the rates were similar between the two age groups (young, 48 ± 14%; aged, 47 ± 25%; mean ± SD; Figure 3B; Supplementary Table S5). Fertilized oocytes were cultured in KSOM(aa) medium and were scored at consecutive stages until the blastocyst stage (96 hours), setting the 2-cell stage as the reference (100%). Within the resolution of our experiment, the rates of cleavage support that embryos derived from maternally aged oocytes were not inferior to those from younger females at any preimplantation stage, as exemplified by blastocyst rates of 73 ± 12% (aged) vs 64 ± 16% (young) (mean ± SD; Figure 3B; Supplementary Table S5). The non-inferiority of maternally aged oocytes, observed after fertilization, was the case also when the developmental stimulus was parthenogenetic activation, which allows to exclude the spermatozoon and thereby isolate the contribution of the oocyte. Parthenogenetically activated, maternally aged oocytes developed to blastocyst at a rate of 80 ± 11% compared to 68 ± 26% of the young counterparts (mean ± SD; N=143 and 237 starting oocytes in three replicates each, respectively; Figure 3B’; Supplementary Table S6).

**Figure 3.**
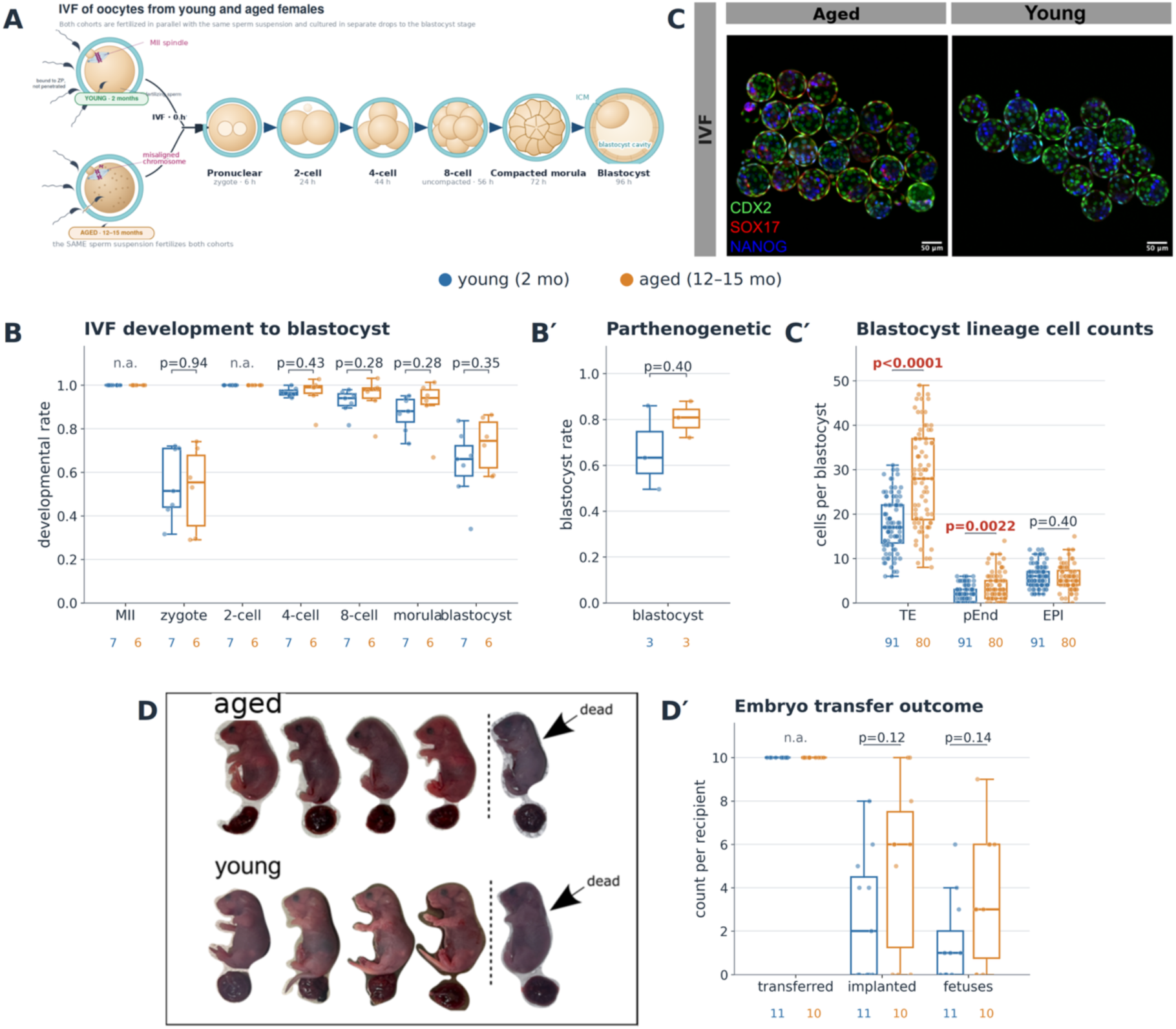
Fertilized embryos do not show the functional deficits expected from maternally aged oocytes. Young = 2 months, aged = 12–15 months (B6C3F1). (A) IVF design: MII oocytes from both cohorts were fertilized in parallel with a single shared CD1 sperm preparation, removing paternal and oviductal confounders. (B) Developmental rate at each stage following IVF, expressed as the fraction of inseminated MII oocytes for the zygote and of 2-cell embryos for all later stages; rates are indistinguishable between age groups throughout (n = 7 young, 6 aged replicates). (B’) Blastocyst rate following parthenogenesis, expressed as the fraction of activated oocytes that progressed to blastocyst; rates are indistinguishable between age groups (n = 3 young, 3 aged replicates). (C) Confocal maximum projections of blastocysts immunostained for CDX2 (trophectoderm), SOX17 (primitive endoderm) and NANOG (epiblast); aged on the left, young on the right. Scale bar = 50 µm. (Cʹ) Cells per blastocyst positive for each marker: aged oocytes yield more trophectoderm and primitive-endoderm cells, with the epiblast unchanged (n = 91 young, 80 aged blastocysts). (D) Embryo transfer: ten embryos were transferred per pseudo-pregnant CD1 females and assessed by Caesarean section at E18.5. (Dʹ) Implantation sites and fetuses per recipient (transferred embryos shown for reference); neither differs significantly, though point estimates trend higher in the aged-donor group (means: implantations 5.1 vs 2.6, p = 0.12; fetuses 3.6 vs 1.5, p = 0.14; n = 11 young, 10 aged recipients). Box plots overlaid with individual data points show the interquartile range, the central line the median and the whiskers the data range excluding outliers. Dots represent the individual replicates (B), blastocysts (B’, C’), recipients (D’); p values by Wilcoxon rank-sum test. Abbreviations: IVF, *in vitro* fertilization; TE, trophectoderm; pEnd, primitive endoderm; EPI, epiblast; Y, young; Ag, aged. **Alt text.** Blastocysts from maternally aged oocytes develop normally. Panel A, scheme of parallel IVF of young and aged oocytes with shared sperm; B, plot of cleavage rates that overlap between age groups at every stage; C, confocal images of blastocysts immunostained for the three lineages, with Cʹ cell-count plots showing more trophectoderm and primitive-endoderm cells in aged embryos; D, embryo-transfer scheme with Dʹ plots of implantation sites and fetuses per recipient.

At the blastocyst stage, the IVF embryos were subjected to differential immunostaining to identify the three cell compartments: trophectoderm (CDX2-positive), primitive endoderm (SOX17-positive), and epiblast (NANOG-positive) (Figure 3C). Blastocysts derived from maternally aged oocytes had more trophectoderm and primitive endoderm cells than their younger counterparts, while the epiblast was unchanged (Figure 3C‘; Supplementary Table S7). However, more cells do not necessarily mean fitter embryos. Assessing fitness would require the examination of further development, which we pursued by transferring embryos in groups of 10 into oviducts of young of pseudo-pregnant recipients (10 and 11 transfers for aged and young, respectively; Supplementary Table S8). Of the 36 and 16 fetuses retrieved in total on developmental day 18.5 after Caesarean section, only one was recovered dead in each group (Figure 3D). In addition to the fetuses, the sites of resorption could still be identified, and by adding the number of resorptions to the number of fetuses we estimated the implantation rate (Figure 3D’). The point estimates trend higher, not lower, in the aged-donor group (implantation 5.1 vs 2.6; fetuses 3.6 vs 1.5 per recipient; n = 11 young, 10 aged; p = 0.12 and 0.14), a direction opposite to most *in-vivo* reports (see Introduction).

Collectively, these data document that in spite of the aging hallmarks found in the oocytes, the embryos showed no detectable functional deficits. With either developmental stimulus, the oocytes of older females formed as many blastocysts - if not more - than those of their younger counterparts, and the blastocysts did not exhibit reduced cell numbers. Past the blastocyst stage, embryo transfer results excluded a large difference of performance between young and aged groups, although it must be acknowledged that minor differences cannot be ruled out, given the size of the embryo transfer groups.

### Fertilization, not mere embryogenesis, enables the directional redistribution of the embryonic proteome

There is no simple way to explain the non-inferior performance of preimplantative mouse embryos originated from maternally aged oocytes fertilized *in vitro*, other than through: 1) counterselection of embryos originated from poor-quality oocytes, or 2) compensatory effects at the gene expression level. Our experimental design cannot exclude counter-selection during preimplantation development, because only oocytes that were fertilized and that made it to the blastocyst stage were effectively considered (survivor bias). However, fertilization rates were similar for young (48%) and aged (47%) oocytes (Figure 3B), after which attrition of compromised embryos should manifest as a lower proportion of fertilized oocytes reaching the blastocyst stage in the aged group. As no such attrition was observed at any preimplantation stage (Figure 3B), counter-selection acting on blastocyst formation is deemed unlikely.

The remaining possibility of compensation prompted us to examine the quality of embryos as measured by global gene expression. Since the aged blastocyst proteome has not been examined before, we chose to confirm the transcriptome first, so as to secure a reproducible comparison before venturing into the unprecedented proteome analysis. Consistent with the published reports (see Introduction), IVF blastocysts showed a predominance of downregulated transcripts (DOWN in aged, n = 6598; STABLE, n = 2369; UP in aged, n = 4008; Supplementary Figure S2, Supplementary Table S9). As for the oocyte proteomes, these are directional assignments rather than a differential-expression analysis. The GO CC terms of these transcripts overlapped substantially with those of the aged oocyte proteome, five of the ten top terms being shared, including ribosome and spliceosomal complex (Supplementary Figure S2, Supplementary Table S9). The picture changed in the blastocyst proteome. Given the smaller number of embryos compared to the initial number of oocytes, and the impossibility to amplify proteins, a low-input pipeline was applied that was originally developed for single-cell proteomics (Zhang et al., 2025a; Materials and Methods). Five samples of three blastocysts each were generated for each maternal age group using either of the two embryo production methods (fertilization and parthenogenesis). Of 4550 proteins (range 4146-4438) that were detected overall in the fertilized blastocysts, 3793 were present in all 10 samples and 2622 were shared with the oocytes (Supplementary Table S10). The main set of IVF embryos was flanked with a parthenogenetic counterpart, so as to resolve if compensation hinges on general processes set in motion during development, or on the paternal contribution. Of 5032 proteins (range 3404-4625) that were detected overall in the parthenogenetic blastocysts, 2757 were present in all 10 samples and 2077 were shared with the oocytes (Supplementary Table S11).

We applied k-means (k = 3) clustering on the Z-normalized values of the blastocyst proteins (rows) and hierarchical clustering on the 10 blastocyst samples (columns) (Figure 4A). Hierarchical clustering assigned all the fertilized samples and all but two of the parthenogenetic samples correctly to the two age groups (young, aged). Given the higher cell counts of blastocysts from aged oocytes (Figure 3C‘), one could object that the two age groups resolved simply because different amounts of cellular material were loaded when the number of embryos per sample is the same (three blastocysts each). Two observations argue against this. First, the number of proteins identified per sample was comparable between groups (Supplementary Table S10). Second, the ACTIN protein DIA values were essentially the same in young and aged samples (ratio aged/young 0.97 after IVF and 0.95 after parthenogenesis); since the *Actin* gene expression level scales to the total number of embryonic cells (Giebelhaus et al., 1983; Giebelhaus et al., 1985), the aged/young ratios do not support that loading played a role. K-means clustering returned the proteins that increase, decrease or remain stable during maternal aging (Figure 4A, A’). As in the case of the oocyte proteome, the abundance-based clustering was directionally sound but statistically permissive, and we retained all cluster members for further analysis regardless of statistical significance (n=865 significant of 1121 DOWN in aged, n=158 significant of 1429 STABLE, n=755 significant of 1243 UP in aged, at raw p < 0.05, t-test, after fertilization; n=173 significant of 1067 DOWN in aged, n=13 significant of 602 STABLE, n=109 significant of 1088 UP in aged, at raw p < 0.05, t-test, after parthenogenesis). In contrast to the aging oocyte proteome (Figure 1 E,E’) and to the aging blastocyst transcriptome (Supplementary Figure S2), the major cluster in aging IVF blastocysts was no longer that of the DOWN (depleted) proteins, but that of the STABLE proteins (Figure 4A).

**Figure 4.**
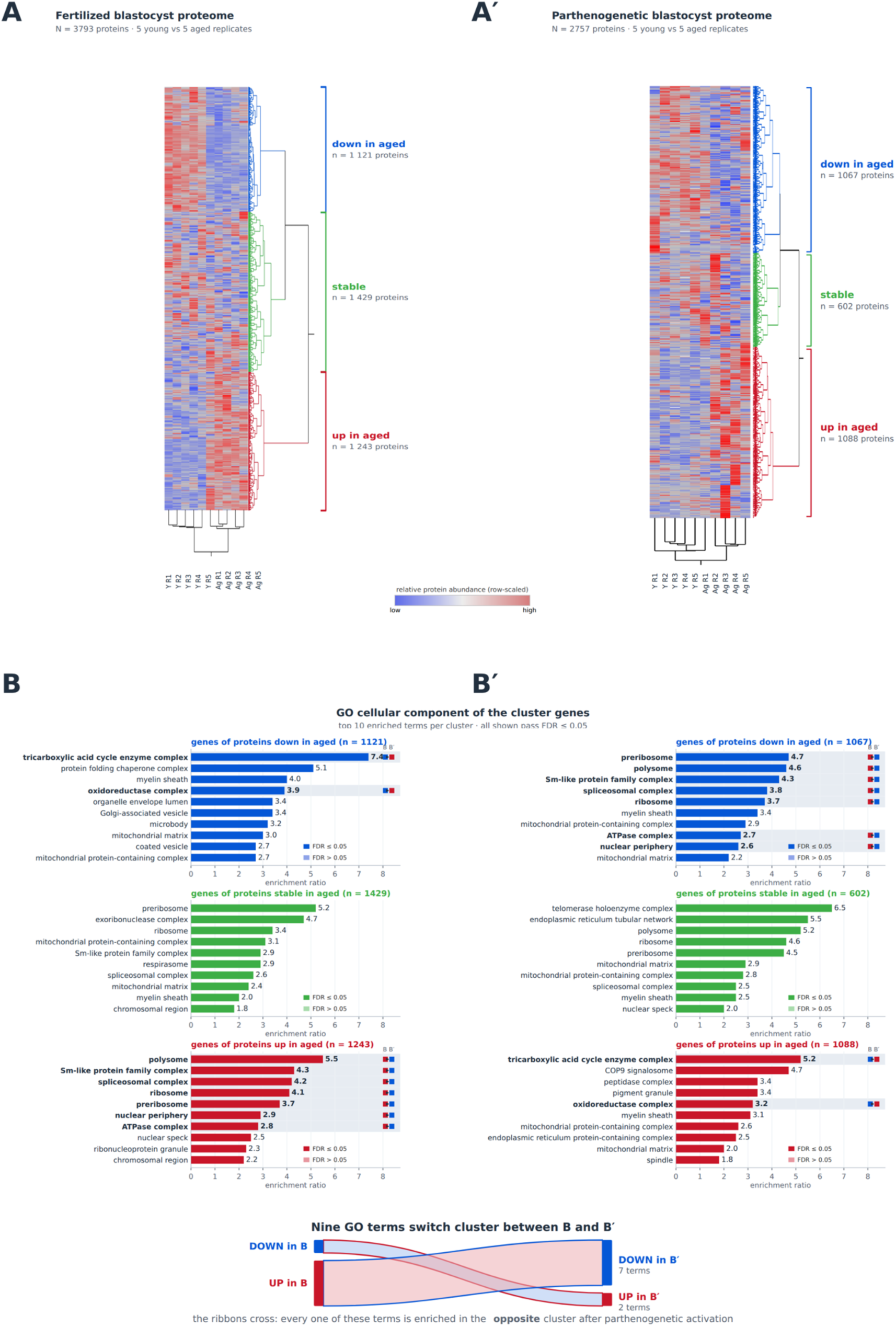
Clustering of the blastocyst proteome by maternal age, with GO (’cellular component’) analysis of the directional clusters. Fertilized (A, B) and parthenogenetic (Aʹ, Bʹ) blastocysts are shown side by side, each from 5 young and 5 aged replicates (young = 2 months, aged = 12–15 months). (A, Aʹ) Heat maps of the proteins quantified in all ten samples of each blastocyst type. Rows are proteins, columns are replicates (Y R1–R5 = young, Ag R1–R5 = aged; R = replicate), and color is row-scaled relative abundance on the shared scale (blue low, red high). Each proteome was clustered independently: columns by hierarchical clustering with Ward linkage, rows partitioned into three clusters by k-means after z-normalisation. (B, Bʹ) Over-representation analysis of each cluster (WebGestalt, ‘cellular component’; reference = whole genome), as enrichment ratio; the ten most enriched terms per cluster are shown and all pass FDR ≤ 0.05. Switching terms are marked three ways — shaded row, bold term and value, and a two-tone chip at the right of the bar giving the cluster color in B then Bʹ. Sankey plot summarizes the nine terms that switch cluster between the two routes of embryogenesis: seven that are UP in fertilized blastocysts are DOWN in parthenogenetic ones, and two that are DOWN in fertilized blastocysts are UP in parthenogenetic ones. Abbreviations: Y, young; Ag, aged. **Alt text.** Heat maps and bar charts of the blastocyst proteome by maternal age. Panels A and Aʹ, side-by-side clustered heat maps of the fertilized and parthenogenetic proteomes, with proteins partitioned by directional change into DOWN (blue), STABLE (green), and UP (red) clusters; panels B and Bʹ, horizontal bar charts of the top-10 most enriched gene-ontology cellular-component terms per cluster, comparing the two embryo types.

We next examined whether the directional protein clusters (Figure 4A,A’) were enriched for specific cellular components (Figure 4B,B’). Over-representation analysis of the directional clusters in the GO CC (reference: protein-coding genome) showed that terms related to ribosomes and mRNA splicing were over-represented among the proteins that went UP with maternal age after fertilization, while terms related to mitochondria were ambivalent (either DOWN or STABLE) (Figure 4B). This is a reversal when one considers how the same terms fared in the original oocytes, particularly clear in the ribosomes and mRNA splicing terms (DOWN with aging in oocytes, UP with aging in blastocysts; DOWN→UP reversal flux). Unlike what was observed after fertilization, the DOWN→UP flux was not pronounced after parthenogenetic activation, and the GO signature of the oocyte DOWN cluster did not become the GO signature of the UP cluster (Figure 4Bʹ). We repeated the over-representation analyses using the detected proteome of the corresponding experiment as reference (Supplementary Figure S1). Although the top-10 GO CC terms of each cluster were only partly retained when using the detected proteome as reference, the salient terms (on which our interpretation rests) were conserved, and remained significant (FDR ≤ 0.05): polysome, ribosome and spliceosomal complex among the proteins that increase with maternal age in fertilized blastocysts, and tricarboxylic acid cycle enzyme complex and oxidoreductase complex among those that decrease. In the parthenogenetic blastocysts, the same categories that characterize the UP cluster after fertilization characterize the DOWN cluster after parthenogenetic activation, although the terms lost significance (FDR > 0.05).

Collectively, these data showed that the GO CC signature of the directional clusters is almost the opposite in the blastocysts after fertilization of what was observed in the oocytes of aging females. For example, terms overrepresented in the DOWN cluster of aging oocytes became those overrepresented in the UP cluster of the blastocysts formed by those aged oocytes. This contrast was not observed after parthenogenesis, leading us to imagine that the key for the compensation may lie in the spermatozoon.

### Fertilization and parthenogenesis redistribute the maternally aged proteome along divergent fluxes

We reasoned that when changes in the abundance of certain protein categories (e.g. ‘ribosome’ or ‘spliceosomal complex’) are observed (Figure 4), these could be the same proteins or distinct proteins that belong to the same GO CC term. We therefore refined the analysis by looking closer at the proteins, in two approaches.

In the first approach (Figure 5A,A’), we examined all detected proteins individually. We plotted the aged/young variations in blastocysts’ proteins against the aged/young variations in oocytes’ proteins, in logarithmic form. The age effects in oocyte and blastocyst correlated significantly negative across the 2622 co-detected proteins after fertilization (Spearman ρ = −0.12, p = 1.1 × 10⁻⁹; Figure 5A). By contrast, after parthenogenesis, the two effects correlated significantly positive across the 2077 co-detected proteins (Spearman ρ = +0.08, p = 5 × 10⁻⁴; Figure 5A’). The two routes of embryogenesis, IVF and parthenogenesis, differed significantly from each other (Fisher’s z = −6.50, p = 8 × 10⁻¹¹). We considered the possibility that these differences could happen by chance, and calculated the probability that random sampling could produce a difference of the magnitude observed (permutation analysis; see Materials and Methods). In the case of the Spearman correlation ρ, the measured value was significantly lower than expected by random re-pairing after fertilization, while it fell in the range expected by random-repairing after parthenogenesis. In the case of the DOWN→UP category, the fertilized value lie above its null (342 observed vs 302 expected; p = 0.001 by permutation), whereas the parthenogenetic value lie below (377 vs 405; p = 0.005; Supplementary Figure S3). Among the detected proteins, four classes already related to previously encountered features (Figure 1E‘, Figure 3C’,Figure 4B) underwent inversion i.e. changed quadrant with age after fertilization (Figure 5A) but not after parthenogenesis (Figure 5A’): spliceosome, ribosome, the SCMC and trophectoderm (Figure 5; Supplementary Table S12). Although all four classes changed quadrant, the seven members of the SCMC class stood out by magnitude of change, as if maternal age would accelerate or intensify the sperm-dependent clearance of the SCMC (Figure 5A; supplementary Figure S4A).

**Figure 5.**
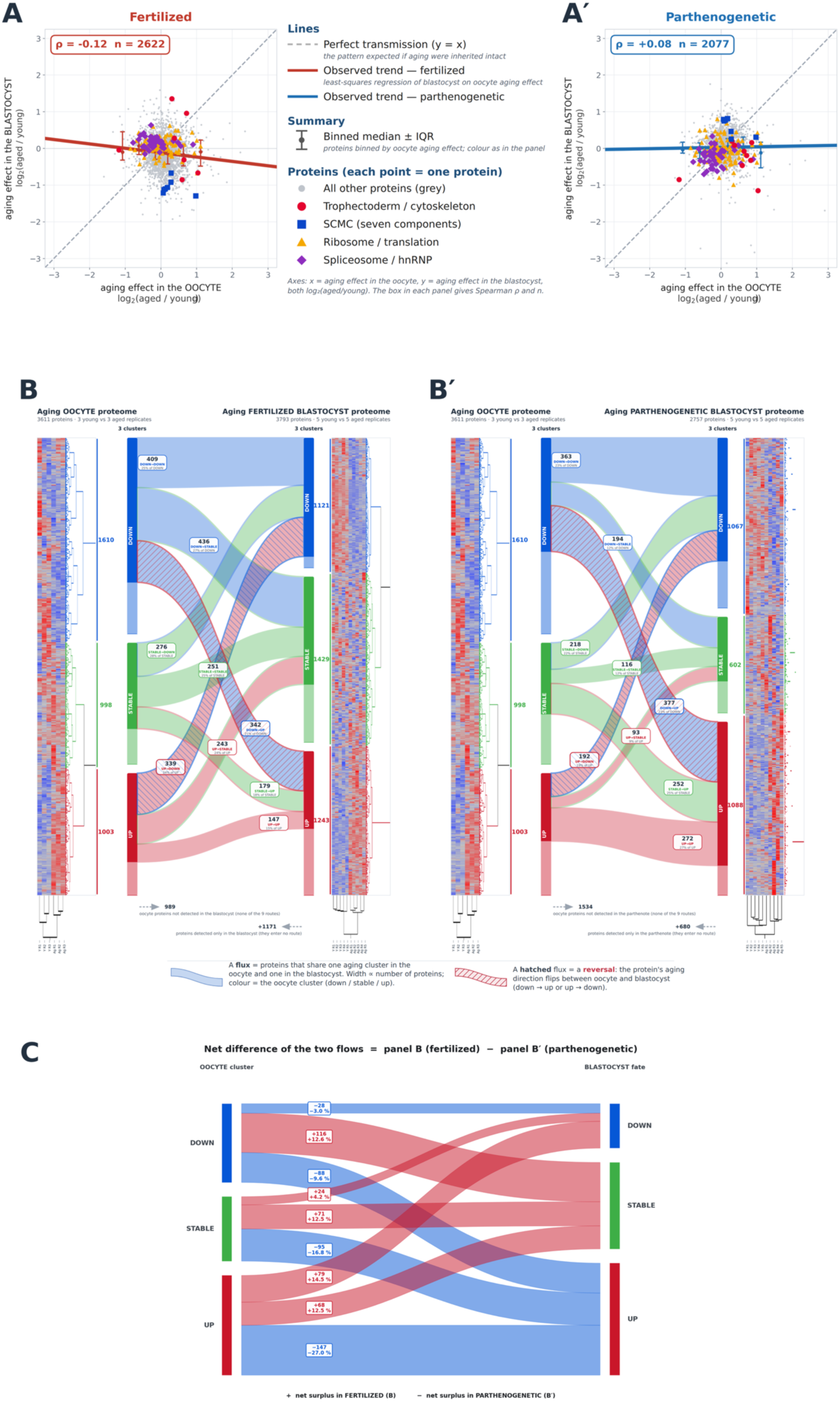
Proteins responding to maternal aging in blastocysts are largely not those that responded in oocytes, more so after fertilization than after parthenogenesis. The aging oocyte proteome (3611 proteins) is traced into fertilized (IVF) blastocysts (A, B) and parthenogenetic blastocysts (Aʹ, Bʹ), and the two embryonic routes are directly compared in a single net-difference plot (C). (A, Aʹ) Protein-level transmission of the maternal-age signature. Each point is one protein quantified at both stages (2622 after fertilization, 2077 after parthenogenetic activation), plotted as its age effect in the oocyte (x) vs in the blastocyst (y), both log₂(aged/young). The dashed diagonal (y = x) signifies perfect transmission; the colored line refers to least-squares regression; whiskered symbols refer to binned medians ± IQR grouped by oocyte age effect. Functional groups are highlighted per the central key; all others grey. Blastocyst values do not follow the diagonal in either route of embryogenesis: the correlation is weakly negative after fertilization (Spearman ρ = −0.12; 1.44% shared variance, i.e. near-absence of transmission) and weakly positive after parthenogenetic activation (ρ = +0.08), the two differing between routes (Fisher’s z = −6.50, p = 8 × 10⁻¹¹). (B, Bʹ) Sankey diagrams of the flux between the three oocyte and three blastocyst clusters (co-detected proteins). Fluxes are shown as ribbons whose fill color is the oocyte cluster of origin, contour color is the blastocyst cluster of destination, width proportional to protein number (counts printed); reversal categories (DOWN→UP, UP→DOWN) hatched. Row percentages do not sum to 100% because part of the proteome is not recovered downstream: 989 (fertilized) and 1534 (parthenogenetic) oocyte proteins were undetected in blastocysts, while 1171 and 680 were blastocyst-only and enter no flux. (C) Net difference between the two redistribution fluxes, obtained by subtracting the parthenogenetic Sankey (Bʹ) from the fertilized Sankey (B). To make the subtraction a true paired comparison, it is computed over the 2031 proteins co-detected in the oocyte and in both blastocyst types (cf. 2622 and 2077 for the individual fluxes). For each of the nine fluxes the traffic shared by the two blastocyst types cancels out and only the net excess remains. Each flux is labelled with the net protein count and the corresponding difference in percentage points of the oocyte cluster; within each oocyte cluster the signed net values sum to zero. The residual pattern is systematic: every net flux entering the STABLE fate is red (proteins directed to STABLE more often after fertilization) and every net flux entering the UP fate is blue (directed to UP more often after parthenogenesis), the largest single term being the retention of UP-cluster proteins in the UP state after parthenogenesis (−147 proteins, −27 percentage points). Cluster color key (node bars, and B, Bʹ): DOWN blue, STABLE green, UP red. Abbreviations: Y, young; Ag, aged. **Alt text.** Five-panel figure tracing 3611 aging-responsive mouse oocyte proteins into fertilized (IVF) and parthenogenetic blastocysts. Panels A and Aʹ are scatter plots of each protein’s aging effect in the oocyte (x-axis) versus the blastocyst (y-axis); the points scatter away from the diagonal in both lineages, showing that the oocyte aging response is largely not carried through to the blastocyst. Panels B and Bʹ are Sankey flux diagrams showing how proteins from the three oocyte clusters (DOWN, STABLE, UP) redistribute among the same three clusters in the blastocyst, with flux width proportional to protein number. Panel C is a net-difference flux diagram (fertilized minus parthenogenetic): red ribbon mark fluxes taken more often after fertilization, blue ribbon taken more often after parthenogenesis. It shows fertilization directing aging-responsive proteins toward the STABLE state and parthenogenesis directing them toward the UP state.

In the second approach (Figure 5B,B’), we examined the proteins not individually but as directional categories (UP, DOWN, STABLE). Since the maternal age effect was scored twice, in the oocyte and in the blastocysts, each time in three directions (UP, DOWN, STABLE), there are a total of 9 ways in which the same proteins can remain in the oocyte’s original directional cluster or switch to a different one: DOWN→DOWN, DOWN→STABLE, DOWN→UP, STABLE→DOWN, STABLE→STABLE, STABLE→UP, UP→DOWN, UP→STABLE, UP→UP (Figure 5B,B’). The redistribution of proteins from oocyte to blastocyst clusters departed significantly from independence in fertilization vs parthenogenesis, indicating that a protein’s aging behavior at the blastocyst stage is not independent of its behavior in the oocyte, and the strength of this dependence differed between developmental stimuli. It was statistically reliable but weak in fertilized embryos (χ² = 31.5, df = 4, p = 2 × 10⁻⁶; Cramér’s V = 0.078; Figure 5B; Supplementary Table S13), and weaker still in parthenogenetic embryos (χ² = 10.7, df = 4, p = 0.031; Cramér’s V = 0.051; Figure 5B’; Supplementary Table S14). To facilitate interpretation, we distilled the difference between the two redistribution patterns by subtracting panel B’ (parthenogenetic) from panel B (fertilized), computed over the 2031 proteins co-detected in the oocyte and in both blastocyst types (Figure 5C). This way, the traffic shared by the two blastocyst types cancels, and only the fluxes on which they diverge remain. The residual pattern is systematic. Read against the blastocyst-fate axis, every net flux entering the STABLE fate is contributed by fertilization (DOWN→STABLE +116, STABLE→STABLE +71, UP→STABLE +68; all +12–13 percentage points of their oocyte cluster), whereas every net flux entering the UP fate is contributed by parthenogenesis (DOWN→UP −88, STABLE→UP −95, UP→UP −147). The single largest net flux is the retention of UP-cluster proteins in the UP state in parthenotes (−147 proteins, −27 percentage points). Figure 5C therefore reduces the whole comparison to one directional statement: from every starting cluster, fertilization drives a net flux of the maternal-age signature toward stability, while parthenogenesis drives it, and holds it, in the up-regulated direction.

Collectively, these data document that the different responses of young and aged oocytes to the developmental stimulus are mediated by the same proteins, which change the directional cluster to which they belong. Although the proteins are conserved, the volumes of their net fluxes differ between fertilization or parthenogenesis.

### SILAC labeling reveals that fertilization remodels the aged maternal proteome through both new synthesis and maternal-protein clearance, only when sperm is present

The fertilization-driven flux toward STABLE (Figure 5C) could reflect either the synthesis of new protein that replaces and dilutes the aged oocytic pool, or the active degradation and clearance of that pool; and the parthenogenetic retention in the UP state could reflect either sustained over-synthesis or a failure to degrade. To distinguish these possibilities, and to determine which arm of proteostasis fertilization engages that parthenogenesis does not, we next measured protein synthesis directly, using the SILAC method (see Materials and Methods). IVF and parthenogenetically activated oocytes were cultured in a modified KSOM, in which arginine and lysine were replaced by isotopic (but non-radioactive) variants (Arg10, Lys8). Newly synthesized proteins are thus marked heavy (H), inherited maternal proteins remain light (L), and the labelling rate is H/(H+L). Since SILAC has not, to our knowledge, been tested for safety when applied to embryo culture, we verified that it would preserve developmental potential, as tested in oocytes of young donors (64.2 ± 14.3 % blastocyst formation, 8 rounds of 360 zygotes each; 34.8 ± 14.9 % development to term, 14 transfers of 8 embryos). With the comfort of these control data, we moved on to the aged oocytes, running five SILAC samples on IVF blastocysts and three on parthenogenetic blastocysts, each including young and aged oocytes. Labelling was globally higher in aged than young blastocysts after fertilization (Figure 6A), but not after parthenogenesis (Figure 6B; Supplementary Table S15) - most markedly among the proteins that increase with age. Grouping proteins by their destination cluster made this explicit. After fertilization, mean labelling rose monotonically across clusters (DOWN 0.36 < STABLE 0.57 < UP 0.68): the proteins elevated with maternal age are predominantly newly made. After parthenogenetic activation the order inverted (UP lowest, 0.45): here the elevated proteins are retained maternal protein, not new synthesis. The extremes of the labelling scale confirmed that the assay was specific for newly synthesized proteins. At one end, SCMC proteins were not labelled at all (rate 0), as expected for purely maternal factors. At the other end, the most heavily labelled proteins were trophectodermal keratins (median 0.88) and translational machinery (ribosomal proteins, median 0.77; proteome median 0.57) - the biosynthetic and cytoskeletal apparatus the embryo rebuilds (Supplementary Figure S4B). That embryonic proteins were labelled and inherited proteins were not labelled (Supplementary Figure S4C), together with the developmental data above, shows the labelling was specific for newly synthesized proteins. The higher labelling rate after fertilization could arise from more synthesis (the H numerator), less retention of maternal protein (the L denominator), or both. We therefore examined each channel separately. In channel H of SILAC, summed DIA intensity was higher in aged than young blastocysts after fertilization (exact rank-sum p = 0.0079, Wilcoxon) but indistinguishable by age after parthenogenesis (p = 1.00 at n = 3; Figure 6C). An orthogonal assay of bulk protein synthesis (OPP incorporation per embryo; Figure 6D) was partly consistent with the summed DIA intensities in channel H: OPP signal intensity changed sign across development after fertilization (aged zygotes incorporated less OPP, aged blastocysts more), whereas after parthenogenetic activation the aged group was higher from the zygote stage onward (five of six stages; Figure 6E,F; Supplementary Table S16). Thus, aged blastocysts incorporated more OPP than young ones. In channel L of SILAC, summed DIA intensity was lower in aged than young blastocysts after fertilization (p = 0.0079) but again indistinguishable after parthenogenesis (p = 1.00; Figure 6G). Both synthesis and clearance therefore contribute, and only when the sperm is present.

**Figure 6.**
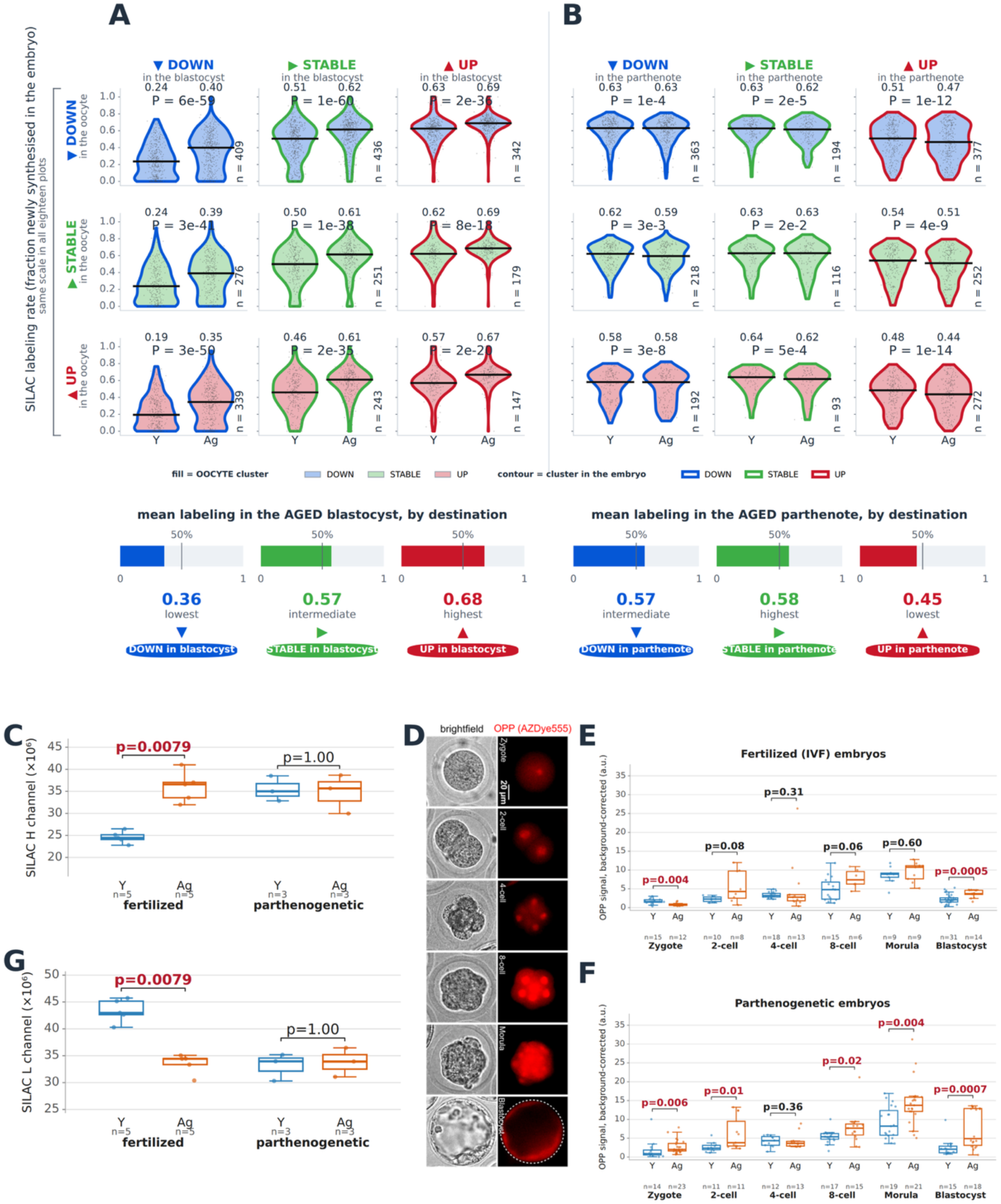
Protein synthesis and maternal-protein clearance in blastocysts from aged oocytes: fertilization versus parthenogenetic activation. **(A, B)** SILAC labeling rate (fraction newly synthesized) in young vs aged blastocysts after fertilization (A; 2622 proteins, 5 vs 5 replicates) and parthenogenetic activation (B; 2077 proteins, 1559 with a value in both ages, 3 vs 3), shown per oocyte → blastocyst flux. Grey dots, individual proteins; black bar, median (printed above); n at lower right, flux size in the contingency table; violins filled by oocyte cluster, contoured by embryo cluster. P values are paired Wilcoxon signed-rank tests across the proteins of each flux. Bars beneath A and B give the mean labeling rate in aged embryos by destination cluster: after fertilization it rises monotonically (DOWN 0.36 < STABLE 0.57 < UP 0.68), so proteins elevated in aged blastocysts are predominantly newly made, whereas after parthenogenetic activation the order inverts (UP lowest, 0.45), indicating retained maternal protein. (C) Newly synthesized protein (SILAC heavy channel): summed DIA intensity of the labelled fraction per sample. Aged blastocysts contain more after fertilization (median 24.5 → 36.6 × 10⁶; exact rank-sum p = 0.0079) but are indistinguishable by age after parthenogenetic activation (35.0 → 35.7 × 10⁶; p = 1.00). n = 5 vs 5 (fertilized), 3 vs 3 (parthenogenetic). (D) OPP incorporation (click chemistry, AZDye555, red = nascent protein) with brightfield beneath, in fertilized embryos at consecutive stages. Scale bar = 20 µm. (E, F) Background-corrected OPP fluorescence per embryo across development after fertilization (E) and parthenogenetic activation (F); background defined by omitting click reagents. After fertilization the age effect changes sign (aged zygotes incorporate less, aged blastocysts more); after parthenogenetic activation the aged group is higher at every significant stage, including the zygote (five of six stages differ). Each dot, one embryo. (G) Inherited maternal protein (SILAC light channel): summed DIA intensity of the unlabelled fraction. Aged blastocysts retain less after fertilization (43.0 → 34.4 × 10⁶; p = 0.0079) but are indistinguishable by age after parthenogenetic activation (34.0 → 33.9 × 10⁶; p = 1.00); n as in C. Together, C and G show that after fertilization, aged oocytes yield blastocysts that both synthesize more protein and clear more maternal protein — the numerator and denominator of the labeling rate in A — with neither channel responding to age after parthenogenetic activation. With three replicates per group the two-sided rank-sum test cannot reach p < 0.05, so the parthenogenetic comparisons establish the absence of a large effect, not of any effect. Throughout: young = blue, aged = orange. Box plots show the interquartile range, the central line the median and the whiskers the data range excluding outliers. Dots represent the individual proteins (A,B) and the individual replicates (C,E,F,G). Significant P (< 0.05) shown in red. Ages compared by Wilcoxon rank-sum test (C, E, F, G). Abbreviations: SILAC, stable isotope labeling with amino acids in cell culture; DIA, data-independent acquisition; OPP, O-propargyl-puromycin; H, heavy; L, light; Y, young; Ag, aged. **Alt text.** Effect of maternal age on embryonic protein synthesis and maternal-protein clearance. Panels A and B, violin plots of SILAC labeling rate per protein by oocyte-to-blastocyst flux, for fertilized and parthenogenetic blastocysts; C, box plots of newly synthesized (heavy-channel) protein; D, micrographs of OPP labeling across development; E and F, box plots of OPP incorporation by stage and age; G, box plots of inherited maternal (light-channel) protein. Labeling and synthesis rise with maternal age only after fertilization.

That both gains depended on the presence of sperm turned our attention to the incoming paternal genome as their trigger, and raised a more speculative question: is the aged oocyte itself better provisioned for the maternal tasks that fertilization demands? In an exploratory, hypothesis-generating analysis *in silico*, we therefore examined whether the oocyte proteins that act directly on the incoming sperm genome are themselves elevated with maternal age. We selected proteins that come to everyone’s mind in this context, i.e. those of the calcium-oscillation apparatus, of sperm-chromatin decondensation and histone deposition, of paternal-genome demethylation, and of ubiquitin–proteasomal clearance. Of 47 such proteins detected (Supplementary Table S17), 21 were more abundant in aged oocytes, 11 less abundant, and 15 unchanged. This runs against the background trend: across the whole oocyte proteome the DOWN cluster was the largest (1610 DOWN, 998 STABLE, 1003 UP), so a random set of 47 would be expected to hold 21 DOWN and 13 UP. The observed excess of UP over DOWN is significant (odds ratio 3.06 vs the whole proteome; Fisher’s exact p = 0.0028; multinomial χ² = 9.88, p = 0.0072). Consistent with being maternally supplied, these proteins were on average inherited rather than newly made (mean labelling rate 0.41 across the 40 with a measurable rate; proteome median 0.57). Four examples illustrate the two earliest steps of paternal-genome handling. Shaping the calcium response to the sperm: the IP₃ receptor ITPR1 (aged/young 1.40; not labelled by SILAC) and the calcium/calmodulin-dependent kinase CAMK2G (1.49; labelling rate 0.19, predominantly inherited), the principal decoder that converts the fertilization calcium oscillations into oocyte activation. Remodelling the sperm chromatin: the nucleoplasmin NPM2 (1.28; labelling rate 0.00, entirely inherited), which exchanges sperm protamines for histones, and the histone chaperone ASF1A (1.81; rate 0.66, largely newly made), which supplies histones for nucleosome assembly on the decondensing paternal DNA.

Taken together, the SILAC and OPP lines of evidence converge on a single, sperm-centered interpretation. Directly, SILAC and OPP show that fertilization - but not parthenogenetic activation - engages both arms of proteostasis in the aged embryo, raising the synthesis of new protein while accelerating the clearance of the inherited maternal pool. Circumstantially, the aged oocyte enters fertilization already enriched for those maternal factors that act first upon the paternal genome, and these are, as expected for stored maternal products, predominantly inherited rather than newly made. Within the statistical power of our experiments, we therefore propose that a maternally-aged proteome is not merely a degraded inheritance to be corrected after fertilization, but is in part provisioned for the earliest steps of paternal-genome handling - offering a candidate basis for the sperm-dependent gains in protein synthesis and maternal-protein clearance that unfold up to blastocyst formation.

## Discussion

The most striking result of this study is that oocytes of aged mice suffer substantial developmental losses when fertilized *in vivo*, but not when fertilized *in vitro*. Once oocytes were separated from the aged fertilization environment, the GO cellular-component categories depleted in the aged oocyte - notably ribosomes and the mRNA-splicing machinery - became the categories enriched among the proteins up-regulated with age in the fertilized blastocyst, a DOWN→UP reversal that is not observed after parthenogenesis. This reversal is not predicted by the prevailing view that oocyte defects are transmitted to the embryo. Two features of our experimental design account for the difference. First, our end point - the direction of a within-stage proteome change - is not the cell number, apoptotic index or transcript abundance on which most prior reports rest. Second, and more importantly, those reports concerned embryos conceived and grown in an aged body, where oocyte age is inseparable from the aged oviduct, the aged uterus (Woods et al., 2017) and, usually, the sperm; our parallel-IVF design removed the paternal variable and the aged tract by construction. Our results therefore complement rather than contradict the literature, not least because no blastocyst-stage proteome stratified by maternal age under equal IVF has been reported before.

Although the maternally aged oocytes showed the expected hallmarks (Introduction; Results), effects in terms of developmental attrition cannot be attributed to the oocyte alone. Bypassing the aged reproductive tract by IVF equalized fertilization rates (47% vs 48%; aged vs young) and blastocyst formation (73% vs 64%), and transfer to young oviducts equalized development to term - so the *in vivo* attrition must arise in part also from the aged maternal environment. Our implantation data after IVF and transfer to young uterus are at variance with the reduced trophectoderm cell number reported for blastocysts of aged mice conceived *in vivo* (Velazquez et al., 2016) and with the recent finding that embryos from aged females implant poorly through a cell-intrinsic increase in trophectoderm contractility (Cavanaugh et al., 2026). In both of those studies the embryos were conceived and developed to blastocyst *in vivo*, within the aged oviduct, whereas ours were produced by IVF and transferred to young oviducts; the trophectoderm phenotype may therefore arise, at least in part, during transit through the aged oviduct rather than being fixed in the oocyte. Consistent with this possibility, our aged-oocyte blastocysts had more - not fewer - trophectoderm cells and no implantation deficit, with epiblast number unchanged. Whether the trophectoderm of IVF-derived aged-oocyte embryos is nonetheless mechanically altered, as reported *in vivo* (Cavanaugh et al., 2026), our cell-count and proteome end points cannot say. The developmental non-inferiority of maternally aged oocytes was also observed without fertilization, that is, in parthenogenesis (80% vs 68% blastocyst formation). Other investigators reported much lower parthenogenetic rates than us for aged oocytes (10.2%), rising to 43.5–50.0% after spindle transfer from aged oocyte to young ooplasm (Yamada and Egli, 2017). Thus, it appears that the mere fertilization of intact oocytes in our study achieved what other investigators obtained by microsurgery, and what dietary NAD⁺ repletion (Bertoldo et al., 2020) or coenzyme Q10 (Ben-Meir et al., 2015) achieved pharmacologically. Although the numerical results differ, these studies converge on a common message: the maternal-age effect on the oocyte is labile, not fixed, and embryos derived from aged oocytes may catch up and become similar to those derived from young oocytes, despite the distinct oocyte qualities. This message echoes our earlier finding that aged and young oocytes yield transcriptionally similar blastocysts after somatic cell nuclear transfer or intracytoplasmic sperm injection, despite divergent oocyte transcriptomes (Esteves et al., 2011). It should be noted that, in this case too (Esteves et al., 2011), the developmental stimulus was applied outside the aged oviduct.

To illuminate the molecular basis of this developmental non-inferiority of maternally aged oocytes when given the developmental stimulus *in vitro*, we compared the age-related changes in blastocysts with those in the precursor oocytes. The blastocyst transcriptome behaved as expected, but the proteome did not: the oocyte signature propagated weakly into parthenogenetic blastocysts (Spearman ρ = +0.08) yet was uncoupled after IVF (ρ = −0.12; Fisher’s z = −6.50, p = 8 × 10⁻¹¹). SILAC qualified this uncoupling in two ways. First, *de novo* protein synthesis (the heavy channel in SILAC) rose after fertilization across all nine redistribution fluxes, indicating a global elevation of embryonic synthesis rather than a targeted repair of the depleted categories. Second, against that raised background there was a modest but reproducible bias toward the depleted proteins: the DOWN→UP flux was over-used after fertilization (342 vs 302 expected; p = 0.001) and under-used after parthenogenesis (377 vs 405; p = 0.005). Restoration was thus achieved by replacement, biased toward the proteins that age had depleted - and it is the biased replacement, not the bias alone, that parthenogenesis failed to reproduce.

Protein synthesis alone did not account for the higher SILAC labelling. OPP, a short pulse of *de novo* synthesis, rose with age after both IVF and parthenogenesis, whereas the integrated SILAC rate rose only after fertilization; because OPP (a 3-h pulse per embryo) and SILAC (a 4-day incubation) measure different quantities, and OPP is sensitive to the higher cell number of aged blastocysts, this divergence is expected. The parsimonious reconciliation is that the inherited maternal pool (the SILAC denominator) was cleared more efficiently after fertilization, which the light-channel intensities confirmed: maternal-protein retention was lower in aged than young blastocysts after IVF but not after parthenogenesis. This agrees with a label-free comparison of fertilized and parthenogenetic embryos from young oocytes, in which certain maternal proteins escaped degradation without sperm (Chen et al., 2022), and with the induction of embryonic autophagy by fertilization but not by parthenogenesis (Tsukamoto et al., 2008). Our data extend this by adding the dimension of maternal age and by distinguishing newly synthesized from inherited protein directly through SILAC.

Why both the gain in synthesis and the loss of maternal protein depend on the spermatozoon is not resolved here, but one candidate concerns how the aged oocyte receives the sperm. The maternal factors acting on the incoming paternal genome - from the calcium-oscillation apparatus of oocyte activation, through the chaperones that decondense sperm chromatin, to the enzymes that clear its packaging - tended to increase with age, suggesting that the aged oocyte enters fertilization well stocked for these earliest, sperm-specific steps. Age alters the fertilization calcium signal itself: aged oocytes sustain normal oscillations after fertilization but reduced ones after parthenogenetic activation (Haverfield et al., 2016), paralleling our pronounced-after-fertilization, muted-after-parthenogenesis proteome remodelling; and components of this machinery were elevated in our aged oocytes, including the IP₃ receptor ITPR1 (1.40×) and its decoder CAMK2G (1.49×). The sperm-chromatin chaperones were likewise elevated — nucleoplasmin NPM2 (1.28×), which exchanges protamines for histones, and ASF1A (1.81×), which supplies histones for the decondensing paternal DNA. We therefore propose, as a testable hypothesis, that maternal age does not merely degrade the oocyte but reconfigures the machinery that receives the spermatozoon, shaping both the activation stimulus and the downstream synthesis-and-clearance programme we observe only when sperm is present.

Despite the precautions of our experimental design that eliminated differences of fertilization environment when comparing the young and aged oocytes side by side, several limitations were nonetheless present in our study. The increased MII aneuploidy of aged oocytes might bias proteome composition in embryos, although the undiminished development to term argues against a large aneuploidy burden, unless aneuploid cells are confined to extraembryonic lineages (Bolton et al., 2016). A single mouse strain was used, and both proteome composition (Pfeiffer et al., 2015) and age-related calcium oscillations (Czajkowska et al., 2020) vary between strains. The IVF rates (fertilized oocytes / inseminated oocytes) were not particularly high (≈50%) in our hands. We realized too late, once the study was already underway, that whilst CD1 males are high performers in *in vivo* embryo production (mating), they are not high performers in IVF (Vergara et al., 1997). Selection at fertilization cannot be excluded, though developmental non-inferiority was previously seen even when all oocytes were fertilized by ICSI (Esteves et al., 2011). The IVF and parthenogenetic routes differed in more than sperm, since parthenotes also received SrCl₂ and latrunculin B, stimuli known to influence development (Ozil et al., 2006). Our aged cohort (12–15 months) was chosen for negligible mortality and thus represents advanced but not terminal reproductive age; a more extreme age might exceed the oocyte’s capacity to be offset. Finally, follow-up ended at birth, leaving open whether latent effects emerge later. Next to these limitations, there is also an observation of uncertain significance: upon scoring the pictures of the maternally young and aged MII oocytes taken side by side immediately after isolation from the cumuli, the aged oocytes had a slightly larger vitellus - opposite to the size recovery reported after nicotinamide mononucleotide treatment of aged females (Bertoldo et al., 2020). We speculate that the size of the vitellus may reflect altered follicular dynamics with age (Zheng et al., 2014) or impaired osmotic regulation after isolation.

To conclude, and bearing in mind our analytical resolution, our statistical power and the limitations, maternally aged oocytes are not inferior to young ones once decoupled from the aged soma and the paternal contribution by IVF. The oocyte’s age signature is not transmitted intact to the blastocyst; it is offset by embryonic protein synthesis and by the clearance of maternal protein - but only when the developmental stimulus is the spermatozoon, not parthenogenesis. This decoupling fits the growing evidence that maternal proteins are exceptionally long-lived, stockpiled in the oocyte and early embryo (Zhang et al., 2023; Harasimov et al., 2024), sequestered in cytoplasmic lattices (Jentoft et al., 2023), and cleared by degradative organelles discharged at maturation and fertilization (Zaffagnini et al., 2024; Satouh et al., 2025). The phenomenon parallels, at the post-transcriptional level, the epigenetic “rejuvenation” by which molecular-clock age falls during early embryogenesis before rising again (Kerepesi et al., 2021), and it reconciles the view that rejuvenation is seeded in the ovary before fertilization (Chiavellini and Sebastiano, 2026) with the view that the epigenetic “age zero” lies at the start of embryogenesis (Kerepesi et al., 2021): the machinery that offsets the maternal-age signature resides in the oocyte but requires fertilization to be engaged. If a comparable offset operates in humans, it would caution against attributing poor outcomes in older women to the oocyte alone and would direct attention also to the peri-fertilization environment - an extrapolation that needs to be tempered by species differences and by our reliance on IVF and culture. The human experiment that would most closely match our mouse design - transferring embryos derived from aged women’s oocytes into young, to test whether a young environment offsets oocyte age - is not performed, and would in any case be confounded by the oocyte aneuploidy that rises with age. What is done clinically is the converse: embryos derived from young-donor oocytes are transferred to aged recipients. In this setting, outcome tracks oocyte age rather than uterine age, with young-donor oocytes largely restoring implantation and pregnancy in older women (Navot et al., 1991) - evidence that the aged uterus is comparatively permissive, as our own transfers to young recipients also imply. Because donation is performed by IVF, however, it bypasses the oviduct and so tests only the oocyte against the uterus, leaving untested the peri-fertilization compartment that our in vivo data implicate. Moreover, the residual age effect that donation cannot rescue is dominated by aneuploidy - the one defect our data, too, place beyond the reach of fertilization. The human evidence thus converges with ours: the rest of the aged oocyte’s molecular legacy is labile rather than fixed.

## Abbreviations

ARRIVE: Animal Research: Reporting of *In Vivo* Experiments
BSA: bovine serum albumin
CC: cellular component (GO analysis)
DIA: data-independent acquisition
FDR: false discovery rate
GO: gene ontology
hCG: human chorionic gonadotrophin
HCZB: Hepes-buffered Chatot, Ziomek and Bavister medium
H&E: hematoxylin and eosin
IF: immunofluorescence
IVF: *in vitro* fertilization
KSOM(aa): potassium simplex optimization medium with amino acids
LC-MS/MS: liquid chromatography with tandem mass spectrometry
MII: metaphase II
OPP: O-propargyl-puromycin
pEnd: primitive endoderm
PMSG: pregnant mare’s serum gonadotrophin
SCMC: subcortical maternal complex
SILAC: stable isotopic labeling by amino acids in cell culture
TE: trophectoderm
WebGestalt: WEB-based GEne SeT AnaLysis Toolkit.

## Statements

### Data availability statement

All data supporting the findings of this study are available in public repositories as listed below and in the Supplementary Materials. The mass spectrometry proteomics data are deposited in the ProteomeXchange Consortium via the PRIDE partner repository under accession numbers PXD076504 (maternally aged mouse oocytes), PXD076689 (fertilized blastocysts), and PXD082343 (parthenogenetic blastocysts). The RNA-seq data are deposited in the ArrayExpress collection in BioStudies under accession numbers E-MTAB-16116. Supplementary Figures 1-4 and Supplementary Tables 1-17 are deposited in FigShare: https://figshare.com/s/8e9cae05c1d9f7d2c9d6

### Ethics statement

This study did not involve human participants, human data or human tissues. Mice were used for experiments according to two licenses issued by the Landesamt für Natur, Umwelt und Verbraucherschutz (LANUV) recently renamed as Landesamt für Verbraucherschutz und Ernährung NRW (LAVE) of the State of North Rhine-Westphalia, Germany (license numbers LANUV AZ.81-02.04.2022.A376 “In-vitro und in-vivo-Markierung der Proteinsynthese in Mausembryonen”, LAVE AZ.2026-273 „Einfluss des maternalen Alters auf die Genexpression von Embryonen im Mausmodell“), in accordance with the procedures laid down in the European Directive 2010/63/EU. Under this regulatory framework, which in Germany is implemented through the Tierschutzgesetz (German Animal Welfare Act) and its subordinate Tierschutz-Versuchstierverordnung, the ethical review and authorization of animal experiments is the legal responsibility of the competent governmental authority—here the LAVE—rather than of an institutional review board (IRB) or an institutional animal care and use committee (IACUC). The licenses cited above therefore constitute the formal ethics approvals for this study and are the European/German equivalent of, and substitutes for, IRB or IACUC approval. The ARRIVE guidelines were observed to the extent applicable. Sample size planning was not possible, given the constraints imposed by the advanced maternal age. The operators were not blind to the age groups, however, the operator who collected the oocytes was distinct from the operator who processed the oocytes. All oocytes were inseminated or activated regardless of morphology.

## Author contributions

According to CRediT (Contributor Roles Taxonomy; https://credit.niso.org) M.B. conceived the study (Conceptualization), made the culture media, performed the embryo culture and embryo transfers (Investigation), analyzed the results (Formal Analysis), drew the figures (Visualization) and wrote the initial draft (Writing – original draft). T.N. helped with the oocyte collection and performed the IF imaging (Investigation). J.W. helped with the oocyte collection and embryo transfers, and performed the OPP assay (Investigation). A.B. and S.W. performed the mass spectrometry analysis (Investigation) under the supervision of H.C.A.D. (Supervision). M.B., H.C.A.D. and G.F. revised the subsequent versions of the manuscript (Writing – review & editing). All authors critically reviewed the manuscript and approved the final version, and agree to be accountable for all aspects of the work.

## Acknowledgments

The authors express gratitude for scientific environment and infrastructural support to the Max Planck Institute for Molecular Biomedicine. As part of infrastructural support, we experienced outstanding support from the mouse housing facility, ensuring a dependable supply of mice needed to collect oocytes and embryos. The RNA-seq analysis was outsourced to the Core Genomic Facility of the Faculty of Medicine of the University of Münster (Anika Witten, Andreas Huge). Part of the results (OPP) presented here were featured in the Master thesis of Julina Wiese.

## Generative AI statement

Claude (Anthropic) was used to assemble the figures from the individual original items either supplied as photographs or generated in JMP, Excel, PowerPoint, Webgestalt. Claude was also used to generate the schematic drawings of embryos, mice and ovaries in Figures 1, 2 and 3, and the Sankey plots in Figures 4 and 5.

## Supplementary Material

The Supplementary Material for this article can be found online at https://figshare.com/s/8e9cae05c1d9f7d2c9d6

**Supplementary Figure S1.** GO cellular-component over-representation analysis repeated with the experiment-detected proteome as the reference set (instead of the whole protein-coding genome), for each proteome (oocyte, fertilized blastocyst, parthenogenetic blastocyst) and each directional cluster (DOWN, STABLE, UP in aged).

**Supplementary Figure S2.** Transcriptome of IVF blastocysts from young (2-month) and aged (12–15-month) B6C3F1 mice: two-way clustering and GO cellular-component over-representation of the directional clusters.

**Supplementary Figure S3.** Two independent tests of whether the oocyte aging signature is reversed in fertilized and parthenogenetic blastocysts: protein-level correlation and permutation of the reversal flux.

**Supplementary Figure S4.** Maternal age accelerates the sperm-dependent clearance of the subcortical maternal complex (SCMC) relative to other abundant embryonic proteins: per-gene maternal-age effect, position within the age-effect distribution, and SILAC labelling.

**Supplementary Table S1.** Body weights, ovary weights, clutch sizes (oocytes ovulated per mouse) of young and aged mice used in this study

**Supplementary Table S2.** Oocyte diameters (vitellus)

**Supplementary Table S3.** Proteome of young and aged oocytes (PXD076504)

**Supplementary Table S4.** Implantations (E7) after mating

**Supplementary Table S5.** Preimplantation rates after IVF

**Supplementary Table S6.** Blastocyst rates after parthenogenesis

**Supplementary Table S7.** Cell lineage allocation in IVF blastocysts

**Supplementary Table S8.** Embryo transfer results after IVF

**Supplementary Table S9.** Transcriptomes of blastocysts formed by young and aged oocytes (E-MTAB-16116)

**Supplementary Table S10.** Proteomes of IVF blastocysts formed by young and aged oocytes (PXD076689)

**Supplementary Table S11.** Proteomes of parthenogenetic blastocysts formed by young and aged oocytes (PXD082343)

**Supplementary Table S12.** Proteins that underwent inversion i.e. changed quadrant with age after fertilization

**Supplementary Table S13.** Redistribution of proteins from oocyte to blastocyst clusters after IVF

**Supplementary Table S14.** Redistribution of proteins from oocyte to blastocyst clusters after parthenogenesis

**Supplementary Table S15.** Labelling (SILAC) in aged vs young blastocysts after fertilization and after parthenogenesis

**Supplementary Table S16.** Labeling of nascent protein synthesis with OPP assay

**Supplementary Table S17.** Oocyte proteins that act directly on the incoming sperm genome, during maternal aging

