## Supplementary figures and tables for "Fertilization offsets the maternal age effect on mouse oocytes provided the oviduct is not itself aged"

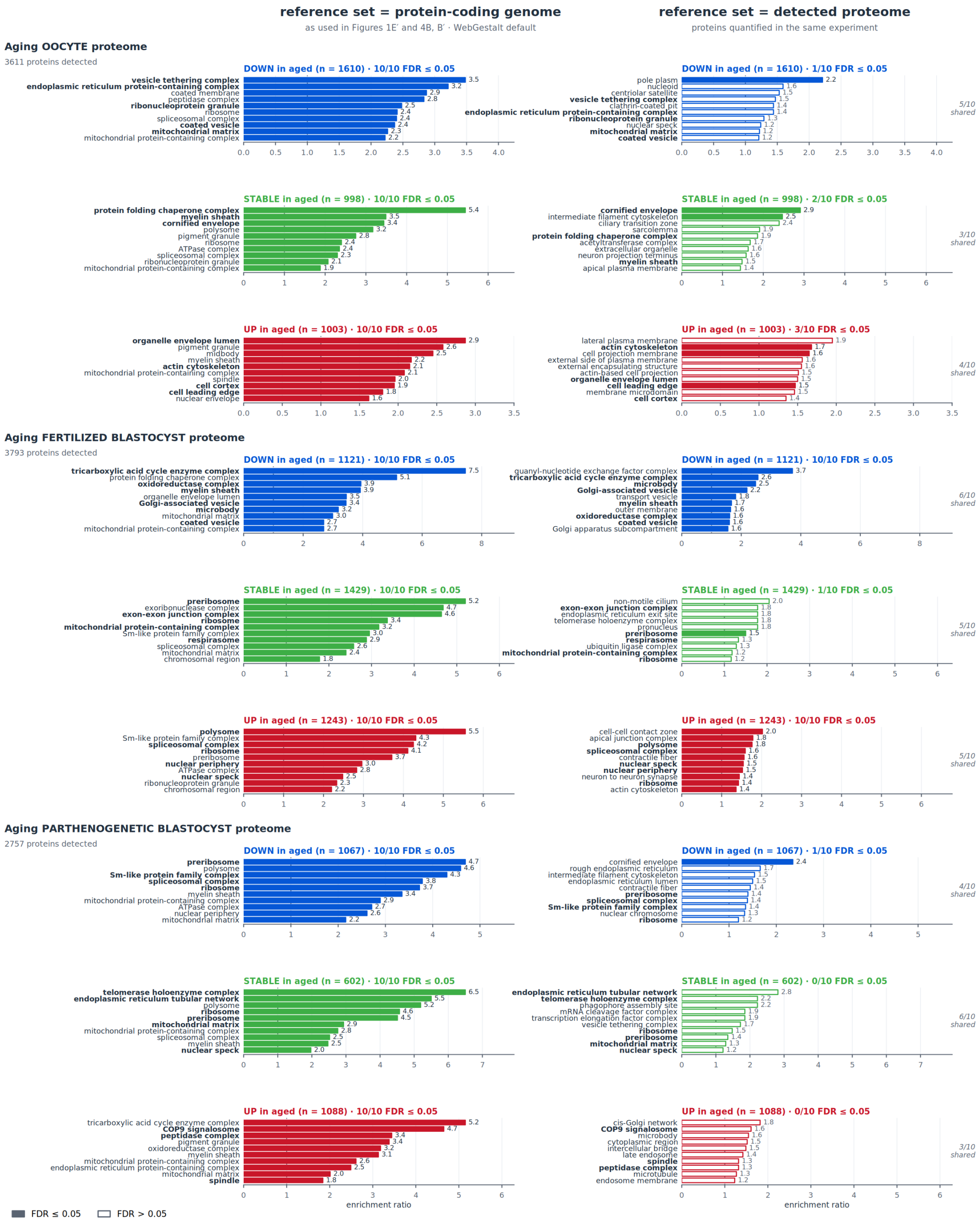


**Supplementary Figure S1. GO cellular-component over-representation analysis repeated with two reference sets.** For each proteome (oocyte, fertilized blastocyst, parthenogenetic blastocyst) and each directional cluster (DOWN, STABLE, UP in aged), the ten most enriched terms are shown against the protein-coding genome (left) and against the proteins detected in the same experiment (right). Within each row the two panels share an x-axis, so the reduction in enrichment ratio is directly comparable. Filled bars pass FDR ≤ 0.05, open bars do not; the count is given in each panel heading. Terms recovered with both reference sets are set in bold, and the number shared is given at the right of each row. Color key as in Figure 4: DOWN, blue; STABLE, green; UP, red.

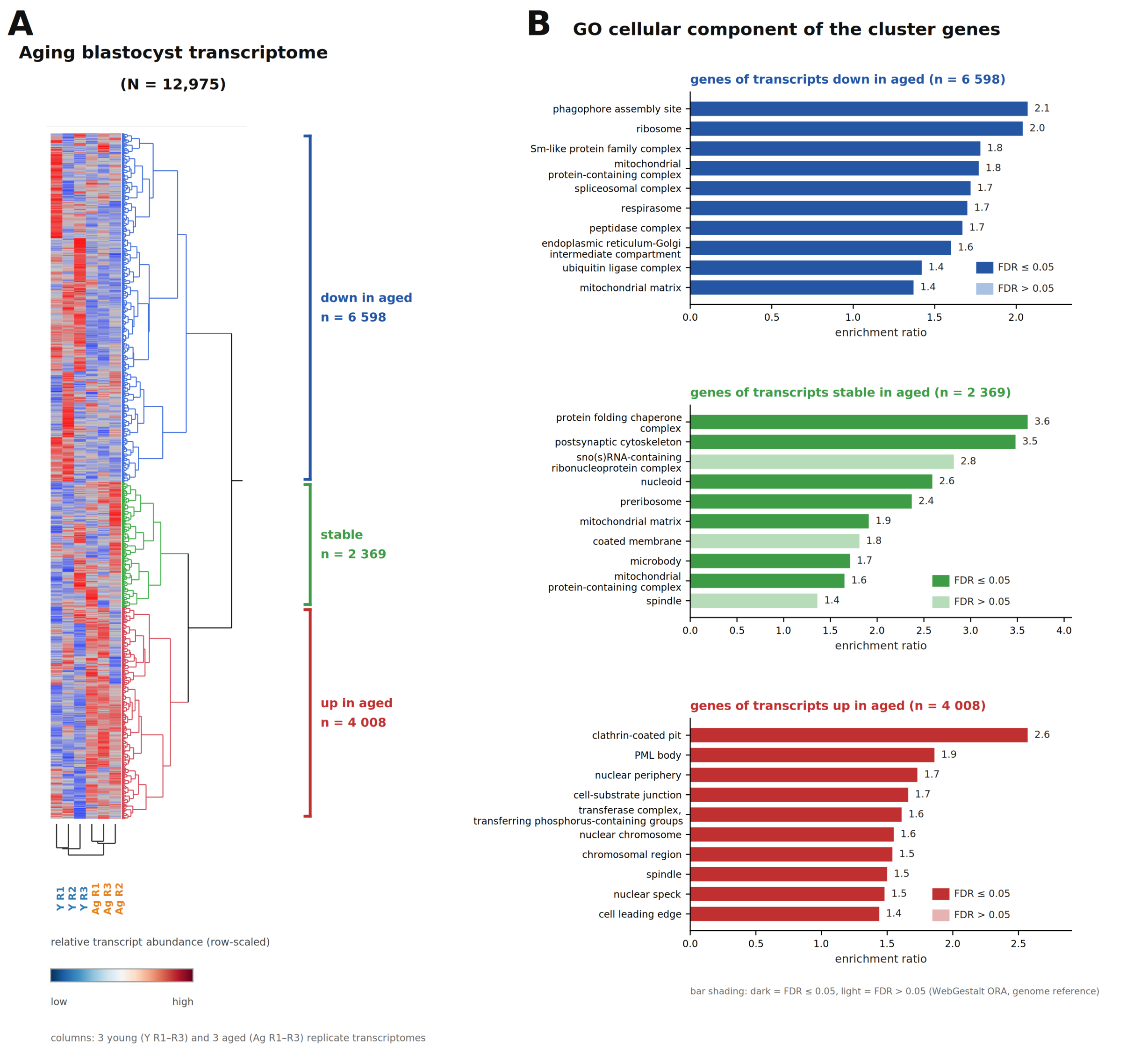


**Supplementary Figure S2. Transcriptome of IVF blastocysts obtained from young (2-month) and aged (12–15-month) B6C3F1 mice.** (A) Two-way clustering of the blastocyst transcriptome (3 young, 3 aged replicates): transcripts quantified in all six samples were z-normalized and grouped by k-means (k = 3, rows); samples (columns) were clustered hierarchically and assigned correctly by age. As in Figures 1E and 4A, cluster membership is a directional assignment. Color = row-scaled relative abundance (blue low, red high). (B) Over-representation analysis of each cluster (WebGestalt, cellular component; reference = whole genome), as enrichment ratio. As in the case of oocytes (Figures 1E, E'), the largest cluster consists of downregulated gene products, and the overrepresented GO CC terms for these overlap with those of the oocytes.

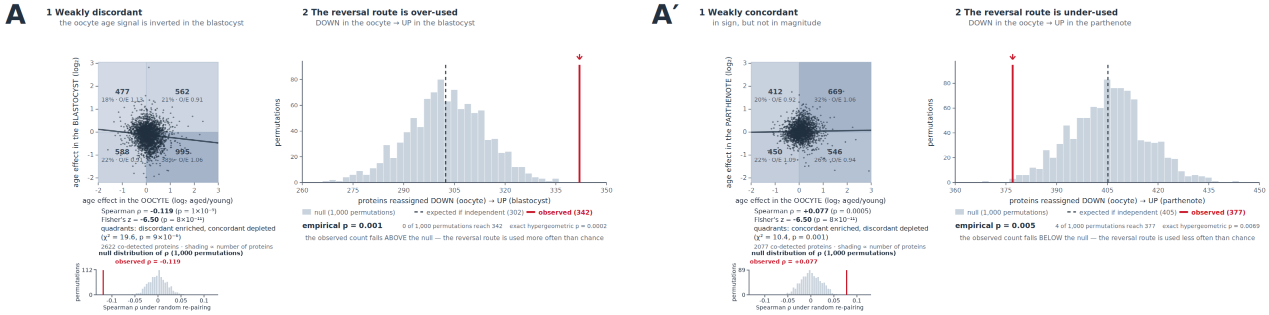


**Supplementary Figure S3. Two independent tests of whether the oocyte ageing signal is reversed in fertilized (A) and parthenogenetic (A′) blastocysts.** Left of each panel, protein-level correlation: each point is one co-detected protein (n = 2622 fertilized, 2077 parthenogenetic), plotted as its maternal-age effect in the oocyte (x) against the same quantity in the blastocyst (y), both $\log_2$(aged/young). The grey line is the least-squares fit; the inset below gives the null distribution of Spearman ρ under 1000 random re-pairings, with the observed value in red. Quadrants are shaded in proportion to the proteins they contain and annotated with that number, its percentage, and the observed/expected ratio (expectation = product of the two marginal distributions). After fertilization the correlation is negative and the discordant quadrants are enriched, i.e. the largely not transmitted ($\rho = -0.12$, $p < 0.001$; quadrant $\chi^2 = 19.6$, $p = 9 \times 10^{-6}$); after parthenogenetic activation it is null to slightly positive ($\rho = +0.08$, $p = 0.0005$). Right of each panel, permutation test of the reversal flux: the number of proteins taking the flux DOWN (oocyte) → UP (blastocyst) under 1000 random permutations of the blastocyst cluster labels, a procedure that preserves both sets of cluster sizes exactly. Grey bars, null distribution; dashed line, expectation under independence; red line, observed value. The reversal flux is over-used after fertilization (342 vs 302 expected, empirical p = 0.001) and under-used after parthenogenetic activation (377 vs 405 expected, empirical p = 0.005).

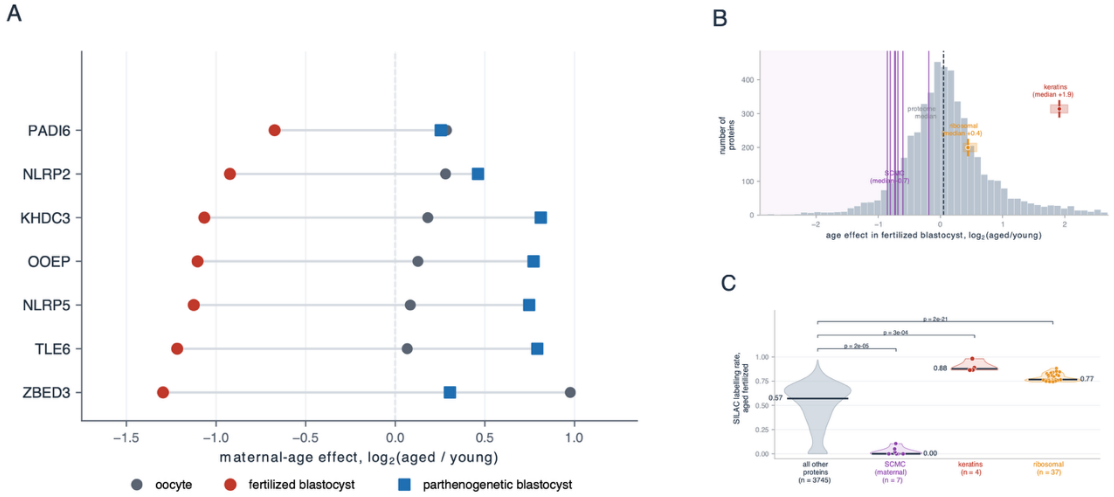


**Supplementary Figure S4. Maternal age accelerates the sperm-dependent clearance of the subcortical maternal complex, compared to other, abundant embryonic proteins**. (A) Per-gene maternal-age effect in oocytes, fertilized blastocysts and parthenogenetic blastocysts for the seven detected SCMC proteins (NLRP5, KHDC3, OOEP, TLE6, PADI6, ZBED3, NLRP2, reviewed in (Bebbere et al., 2021)). All seven are depleted after fertilization and elevated after parthenogenetic activation (paired Wilcoxon p = 0.0156). (B) Position of the seven SCMC members within the distribution of maternal-age effects in fertilized blastocysts (DIA dataset, n = 3793 proteins), compared with ribosomal proteins (n = 37) and keratins (n = 4). (C) SILAC labeling rate in aged fertilized blastocysts. SCMC members are inherited rather than newly synthesized (median 0.00); keratins (KRT19, KRT18, KRT8, KRT5) are almost completely labelled (median 0.88; KRT19 0.98) and ribosomal proteins substantially labelled (median 0.77), against a proteome median of 0.57, showing that they undergo replacement from oocyte to blastocyst (RPS26, RPL10L, RPL7L1, RPL22L1, RPS29, RPS28, RPL39, RPS21, RPS16, RPS14, RPL27, RPS15A, RPL13, RPL18, RPS18, RPS6, RPS3A, RPL19, RPL26, RPL8, RPL4, RPS20, RPL13A, RPS13, RPS12, RPS5, RPS9, RPS19BP1, RPS8, RPS15, RPL22, RPL15, RPS25, RPL7, RPL27A, RPS23, RPSA).

### Supplementary Tables

Supplementary Tables 1–17 are provided as separate data files and are deposited, together with the underlying source data, in the figshare repository:

https://figshare.com/s/8e9cae05c1d9f7d2c9d6

**Supplementary Table S1.** Body weights, ovary weights, clutch sizes (oocytes ovulated per mouse) of young and aged mice used in this study
**Supplementary Table S2.** Oocyte diameters (vitellus)
**Supplementary Table S3.** Proteome of young and aged oocytes (PXD076504)
**Supplementary Table S4.** Implantations (E7) after mating
**Supplementary Table S5.** Preimplantation rates after IVF
**Supplementary Table S6.** Blastocyst rates after parthenogenesis
**Supplementary Table S7.** Cell lineage allocation in IVF blastocysts
**Supplementary Table S8.** Embryo transfer results after IVF
**Supplementary Table S9.** Transcriptomes of blastocysts formed by young and aged oocytes (E-MTAB-16116)
**Supplementary Table S10.** Proteomes of IVF blastocysts formed by young and aged oocytes (PXD076689)
**Supplementary Table S11.** Proteomes of parthenogenetic blastocysts formed by young and aged oocytes (PXD082343)
**Supplementary Table S12.** Proteins that underwent inversion i.e. changed quadrant with age after fertilization
**Supplementary Table S13.** Redistribution of proteins from oocyte to blastocyst clusters after IVF
**Supplementary Table S14.** Redistribution of proteins from oocyte to blastocyst clusters after parthenogenesis
**Supplementary Table S15.** Labelling (SILAC) in aged vs young blastocysts after fertilization and after parthenogenesis
**Supplementary Table S16.** Labeling of nascent protein synthesis with OPP assay
**Supplementary Table S17.** Oocyte proteins that act directly on the incoming sperm genome, during maternal aging

The mass spectrometry proteomics data are available via ProteomeXchange/PRIDE under accession numbers PXD076504 (maternally aged mouse oocytes), PXD076689 (fertilized blastocysts) and PXD082343 (parthenogenetic blastocysts); the RNA-seq data are available in BioStudies/ArrayExpress under accession number E-MTAB-16116.
